# Improved cortical surface reconstruction using sub-millimeter resolution MPRAGE by image denoising

**DOI:** 10.1101/2020.09.20.304824

**Authors:** Qiyuan Tian, Natalia Zaretskaya, Qiuyun Fan, Chanon Ngamsombat, Berkin Bilgic, Jonathan R. Polimeni, Susie Y. Huang

**Author notes:** Correspondence to: Qiyuan Tian, Ph.D., Athinoula A. Martinos Center for Biomedical Imaging, 149 13th Street, Charlestown, MA, 02129, United States.

## Abstract

Automatic cerebral cortical surface reconstruction is a useful tool for cortical anatomy quantification, analysis and visualization. Recently, the Human Connectome Project and several studies have shown the advantages of using T_1_-weighted magnetic resonance (MR) images with sub-millimeter isotropic spatial resolution instead of the standard 1-millimeter isotropic resolution for improved accuracy of cortical surface positioning and thickness estimation. Nonetheless, sub-millimeter resolution images are noisy by nature and require averaging multiple repetitions to increase the signal-to-noise ratio for precisely delineating the cortical boundary. The prolonged acquisition time and potential motion artifacts pose significant barriers to the wide adoption of cortical surface reconstruction at sub-millimeter resolution for a broad range of neuroscientific and clinical applications. We address this challenge by evaluating the cortical surface reconstruction resulting from denoised single-repetition sub-millimeter T_1_-weighted images. We systematically characterized the effects of image denoising on empirical data acquired at 0.6 mm isotropic resolution using three classical denoising methods, including denoising convolutional neural network (DnCNN), block-matching and 4-dimensional filtering (BM4D) and adaptive optimized non-local means (AONLM). The denoised single-repetition images were found to be highly similar to 6-repetition averaged images, with a low whole-brain averaged mean absolute difference of ∼0.016, high whole-brain averaged peak signal-to-noise ratio of ∼33.5 dB and structural similarity index of 0.92, and minimal gray matter–white matter contrast loss (2% to 9%). The whole-brain mean absolute discrepancies in gray–white surface placement, gray–CSF surface placement and cortical thickness estimation were lower than 165 μm, 155 μm and 145 μm—sufficiently accurate for most applications. The denoising performance is equivalent to averaging ∼2.5 repetitions of the data in terms of image similarity, and 1.6–2.2 repetitions in terms of the cortical surface placement accuracy. The scan-rescan precision of the cortical surface positioning and thickness estimation was lower than 170 μm. Our unique dataset and systematic characterization support the use of denoising methods for improved cortical surface reconstruction sub-millimeter resolution.

## Introduction

Cerebral cortical surface reconstruction enables the delineation of the inner and outer boundaries of the highly folded human cerebral cortex from anatomical magnetic resonance imaging (MRI) data using geometric models of the cortex. The input for cortical surface reconstruction is typically a T_1_-weighted structural MR image volume demonstrating high gray matter–white matter and gray matter–cerebrospinal fluid (CSF) contrast. The output of cortical surface reconstruction is a pair of two-dimensional surfaces represented as polyhedral (triangular) surface meshes, which represent the gray matter–white matter interface and the gray matter–CSF interface. Accurate cortical surface reconstruction can be performed automatically using several publicly available software packages^1–9^ and has been routinely used to quantify cortical morphology^10–24^, parcellate cortical areas^25–27^, and visualize and analyze anatomical^28–32^, microstructural^33–36^ and functional^37–42^ signals at various cortical depths and on the folded cortical manifold in a wide range of clinical and neuroscientific applications.

High-quality input T_1_-weighted MR images are essential for generating accurate cortical surfaces. T_1_-weighted MR images acquired with 1-millimeter isotropic spatial resolution have been long used for standard cortical surface reconstruction because the 1-millimeter voxels adequately sample the folded cortical ribbon in adult humans and because this acquisition provides an adequate trade-off between acquisition time, spatial resolution and signal-to-noise ratio (SNR). More recently, the Human Connectome Project (HCP) WU-Minn-Ox Consortium and several studies have shown the advantages of sub-millimeter isotropic spatial resolution for further improving the accuracy of the reconstructed cortical surfaces^5,43–47^. In these studies using sub-millimeter cortical surface reconstruction, the cortical surface positioning and thickness estimation in cortical regions with diminished gray–white contrast on T_1_-weighted MR images exhibit improved accuracy due to reduced partial volume effects on the sub-millimeter resolution images. This effect is especially noticeable in heavily myelinated cortical areas (e.g., primary motor, somatosensory, visual and auditory cortex), where the gray matter signal appears more similar in signal intensity to the white matter, as well as in insular cortex where the superficial white matter in the extreme capsule immediately beneath the insular cortex appears darker due to partial volume effects with the claustrum^43,47^. In general, T_1_-weighted MR images with 0.8-mm isotropic resolution (half of the minimum thickness of the cortex) or higher resolution are recommended by the HCP WU-Minn-Ox Consortium^46^. This new standard was adopted in the healthy adult HCP^44,46^ and the subsequent HCP Lifespan Development and Aging^48–50^. On the surface reconstruction side, several software packages, such as BrainVoyager^51^, CBS Tools^5^ and FreeSurfer^43^ have evolved to process anatomical MRI data at native sub-millimeter resolution.

A significant barrier to the wide adoption of the cortical surface reconstruction at sub-millimeter resolution is the longer scan time. Modern accelerated MRI techniques (e.g., compressed sensing^52^ and Wave-CAIPI^53^) can encode a larger number of k-space data samples at the sub-millimeter acquisition with less noise enhancement and artifact penalties compared to conventional parallel imaging approaches. Nevertheless, sub-millimeter resolution MR images remain limited in image quality by the intrinsically low SNR of smaller voxels. Adequate image quality using such data for cortical surface reconstruction requires averaging data from at least two repetitions of the acquisition, resulting in scan times that last ∼15 minutes (assuming a standard imaging acceleration factor of two) compared to ∼5 minutes scan time for a routine 1-mm isotropic resolution acquisition. In fact, the number of the repetitions required to provide the same SNR scales with the square of the voxel size ratio (e.g., ∼3.81 repetitions of 0.8-mm isotropic resolution data are needed to match the SNR of a single repetition of 1-mm isotropic resolution data, assuming all other parameters are held constant). Such long scans are uncomfortable and may result in increased vulnerability to motion artifacts and reduced scan throughput. Ultra-high field (e.g., 7-Tesla and 9.4-Tesla) MRI scanners and highly-parallelized phased-array radio frequency receive coil head arrays (i.e., 64 channels and above^54^) can significantly boost sensitivity and image SNR, but are currently still expensive and less accessible. Even at ultra-high fields, the SNR of a single repetition of sub-millimeter data may be not sufficient for accurately delineating the cortical surface in the temporal lobes because of signal dropout due to dielectric effects, and more repetitions are needed to enable high-quality whole-brain analysis.

Image denoising provides a feasible alternative strategy to obtain high-quality sub-millimeter anatomical MR images with adequate SNR from a single acquisition. Image denoising aims to recover a high-quality image from its noisy, degraded observation, which is a highly ill-posed inverse problem. Denoising algorithms often rely on prior knowledge as supplementary information to estimate high-quality images. One category of denoising methods incorporate these priors, such as sparseness^52,55–58^ and low rank^59–64^, during the formation of MR images. Another category of denoising methods are designed to be applied to the reconstructed images, with numerous algorithms proposed for processing 2-dimensional images (e.g., total variation denoising^65^, anisotropic diffusion filtering^66,67^, bilateral filtering^68^, non-local means (NLM) filtering^69^, block-matching and 3-dimentional filtering (BM3D)^70^ and K-SVD denoising^71^) and 3-dimensional medical imaging data^72–76^. As a stand-alone processing step, these methods only take in reconstructed images from the MRI scanner as inputs, without any need to intervene in the current MRI workflow, and can be directly incorporated into the existing surface reconstruction software packages.

Convolutional neural networks (CNNs) are advantageous for learning the underlying image priors and resolving the complex high-dimensional mapping from noisy images to noise-free images. Even a simple CNN for denoising (i.e., DnCNN^77^), comprised of stacked convolutional filters paired with rectified linear unit (ReLU) activation functions, achieves superior performance compared to the state-of-the-art BM3D denoising method. It is also relatively easy for CNNs to incorporate redundant information from other imaging modalities (e.g., using MR images to assist removal of noise in positron emission tomography (PET) images^78^) to boost denoising performance. Furthermore, denoising CNNs can be trained without use of noise-free target images in a self-supervised manner while still outperforming the state-of-the-art methods, which is especially useful when the noise-free target images are not available^79,80^. Impressively, CNN-based denoising methods can be executed extremely efficiently (in milliseconds or seconds) once trained. Thus far, CNN-based denoising has been widely applied for various biomedical imaging modalities ranging from fluorescence microscopy^80,81^, optical coherence tomography^82^ to x-ray imaging^83^, x-ray computed tomography^84^, PET^78,85–88^ and MRI^84,89–95^.

In the present study, we leverage multiple available denoising methods for improving cortical surface reconstruction at sub-millimeter resolution and characterize the improvement in the reconstructed surfaces offered by the different denoising approaches using empirical data. Specifically, we reconstruct cortical surfaces using single-repetition 0.6-mm isotropic resolution T_1_-weighted images denoised by classical denoising methods, including BM4D^75^, adaptive optimized NLM (AONLM)^74^ and DnCNN^77^. We systematically quantify the similarity and image contrast of denoised 0.6-mm isotropic T_1_-weighted images to the corresponding high-SNR 0.6-mm isotropic images obtained by averaging 6 repetitions of the data acquired within a single scan session and compare the results to those obtained from 2- to 5-repetition averaged data. We also quantify and compare the quality and smoothness of the reconstructed surfaces as well as the similarity of cortical surface positioning and cortical thickness estimation compared to the ground truth. Finally, we characterize the scan–rescan precision of the cortical surface positioning and cortical thickness estimation derived from the sub-millimeter resolution images denoised by different methods.

## Materials and Methods

### Data acquisitions

T_1_-weighted images with 0.6-mm isotropic resolution of nine healthy subjects (4 for training and validation of DnCNN parameters, 5 for evaluation) were acquired on a whole-body 3-Tesla scanner (MAGNETOM Trio Tim system, Siemens Healthcare, Erlangen, Germany) using the vendor-supplied 32-channel receive coil, body transmit coil, and standard body gradients at the Athinoula A. Martinos Center for Biomedical Imaging of the Massachusetts General Hospital (MGH) with Institutional Review Board approval and written informed consent of the volunteers. The data were acquired using a 3-dimensional multi-echo magnetization-prepared rapid gradient-echo (MEMPRAGE) sequence^96^. A slab-selective oblique-axial acquisition using a 13 ms FOCI adiabatic inversion pulse^97^ and acceleration factor of 2 in the partition (i.e., inner loop) direction was used to minimize the number of partition encoding steps and minimize T_1_ blurring during the inversion recovery^98–100^. Because the profile of the radiofrequency pulse for the slab excitation falls off at the slab boundaries, the signal level is lower toward the superior and inferior brain yielding lower SNR locally in the acquired images (Supplementary Fig. 1a). In order to avoid spatial blurring, partial Fourier imaging was not used in any encoding direction to help preserve the 0.6-mm isotropic resolution. The sequence parameters were: repetition time = 2,510 ms, echo times = 2.88/5.6 ms, inversion time = 1,200 ms, excitation flip angle = 7°, bandwidth = 420 Hz/pixel, echo spacing = 8.4 ms, 224 axial slices with 14.3% slice oversampling, matrix size = 400×304, slice thickness = 0.6 mm, field of view = 240 mm × 182 mm, generalized autocalibrating partial parallel acquisition (GRAPPA) factor = 2, acquisition time = 10.7 minutes per run × 6 separate repetitions.

### Image processing

Image co-registration was performed six times for each subject. Each time, five image volumes were individually co-registered to the remaining one image volume (the reference image volume which was not resampled) using the robust registration method as implemented in the “*mri_robust_template*” function^101^ from the FreeSurfer software^1–3^ (version v6.0, https://surfer.nmr.mgh.harvard.edu), with “*satit*” and “*fixtp*” options and “*inittp*” set to the number of the reference image volume. The reference image volume was not re-sampled to avoid changing the noise characteristics. The co-registered image volumes were gradually added to the reference image volume to create averaged image volumes with progressively increasing SNR (*n*=2, 3, 4, 5, 6), which were compared to the de-noised single-repetition data in order to determine the *efficiency factor*, i.e., the number of image volumes for averaging that would be needed to obtain equivalent image and surface metric values compared to those obtained from denoised single-repetition data. Consequently, 6 pairs of image volumes, each pair consisting of a noisy image volume from the single-repetition data and a high-SNR image volume from the 6-repetition averaged data, were generated for each subject. A total of 24 pairs of image volumes (6 pairs/subject × 4 training subjects) were used as the training and validation data for optimizing the CNN parameters. For evaluation, 30 pairs of image volumes, each pair consisting of a noisy image volume from the single-repetition data (or 2- to 5-repetition averaged data) and a high-SNR image volume from 6-repetition averaged data, were used.

Spatially varying intensity bias maps were estimated using the image volumes from the single-repetition data with the unified segmentation routine^7^ implementation in the Statistical Parametric Mapping software (SPM, https://www.fil.ion.ucl.ac.uk/spm) with a full-width at half-maximum (FWHM) of 30 mm and a sampling distance of 2 mm^102^ (Supplementary Fig. 1b). The estimated bias maps were divided from images from the single-repetition and 2- to 6-repetition averaged data for correcting spatially varying intensity bias.

For improved DnCNN performance and evaluation of image and surface quality compared to those obtained from the 6-repetition averaged data, the images from the 6-repetition data were further non-linearly co-registered to the images from the single-repetition data, as well as from the 2- to 5-repetition averaged data, respectively. The non-linear co-registration slightly adjusted the image alignment locally, which accounted for the subtle non-linear shifts of tissue in the images due to factors such as subject bulk motion and the associated changes in distortion due to changing B0 inhomogeneity and gradient nonlinearity distortion during the acquisition of each repetition of data. The non-linear co-registration was performed using the “*reg_f3d*” function (default parameters, spline interpolation) from the NiftyReg software (https://cmiclab.cs.ucl.ac.uk/mmodat/niftyreg^103,104^.

The local mean values of the image intensity from the co-registered 6-repetition averaged data were also slightly scaled to match those of the corresponding single repetition as well as 2- to 5-repetition averaged data to remove the residual intensity bias for improved DnCNN performance and image and surface similarity evaluation. The local mean image intensity was computed using a 3-dimenstional Gaussian kernel with a standard deviation of 2 voxels (i.e., 1.2 mm).

Brain masks were created from the 6-repetition averaged image volumes by performing the FreeSurfer automatic reconstruction processing for the first five preprocessing steps (FreeSurfer’s “*recon-all*” function with the “-*autorecon1*” option). The binarized brain masks were dilated twice with holes filled using the “*fslmaths*” function from the FMRIB Software Library software^105–107^ (FSL, https://fsl.fmrib.ox.ac.uk).

### BM4D denoising

T_1_-weighted image volumes from the single-repetition data were denoised using the BM4D denoising algorithm^70,75^. BM4D extends the widely known BM3D algorithm designed for 2-dimenstion image data to process 3-dimensional volumetric data. The BM4D method provides superior denoising performance by grouping similar 3-dimensional blocks across the entire volume (both local and non-local) into 4-dimensional data arrays to enhance the sparsity and then performing collaborative filtering. The BM4D denoising was performed assuming Rician noise with an unknown noise standard deviation and was set to estimate the noise standard deviation and perform collaborative Wiener filtering with “modified profile” option using the publicly available MATLAB-based software (https://www.cs.tut.fi/~foi/GCF-BM3D).

### AONLM denoising

T_1_-weighted image volumes from the single-repetition data were also denoised using the AONLM algorithm^73,74^. In AONLM, the restored intensities of a small block are the weighted average of the intensities of all similar blocks within a predefined neighborhood. The 3-dimensional AONLM extends the 2-dimensional NLM filtering by using a block-wise implementation with adaptive smoothing strength based on automatically estimated spatially varying image noise level. The AONLM was performed assuming Rician noise with 3×3×3 block and 7×7×7 search volume^73,74^ using the publicly available MATLAB-based software (https://sites.google.com/site/pierrickcoupe/softwares/denoising-for-medical-imaging/mri-denoising/mri-denoising-software).

### DnCNN denoising

T_1_-weighted image volumes from the single-repetition data were also denoised using DnCNN^77^ (Fig. 1), a pioneering and classical CNN for denoising. The 2-dimensional convolution used for processing 2-dimenstional images was extended to 3-dimensional for MRI to increase the data redundancy from an additional spatial dimension for improved denoising performance and smooth transition between 2-dimensional image slices. DnCNN maps the input image volume to the noise, i.e., the residual volume between the input and output image volume. Residual learning facilitates the DnCNN to only learn the high-frequency information, which boosts CNN performance, accelerates convergence and avoids the vanishing gradient problem^77,108,109^. DnCNN has a simple network architecture comprised of stacked convolutional filters paired with rectified linear unit (ReLU) activation functions and batch normalization functions. The *n*-layer DnCNN consists of a first layer of paired 3D convolution (k kernels of size *d*×*d*×*d*×1 voxels, stride of 1×1×1 voxel) and ReLU activation, *n* − 2 middle layers of 3D convolution (*k* kernels of size *d*×*d*×*d*×*k* voxels, stride of 1×1×1 voxel), batch normalization and ReLU activation, and a last 3D convolutional layer (one kernel of size *d*×*d*×*d*×*k* voxels, stride of 1×1×1 voxel). The output layer excludes ReLU activation so that the output would include both positive and negative values. Parameter value settings of *n*=20, *k*=*64*, *d*=3 were adopted following previous studies^77^ (∼2 million trainable parameters).

**Figure 1.**
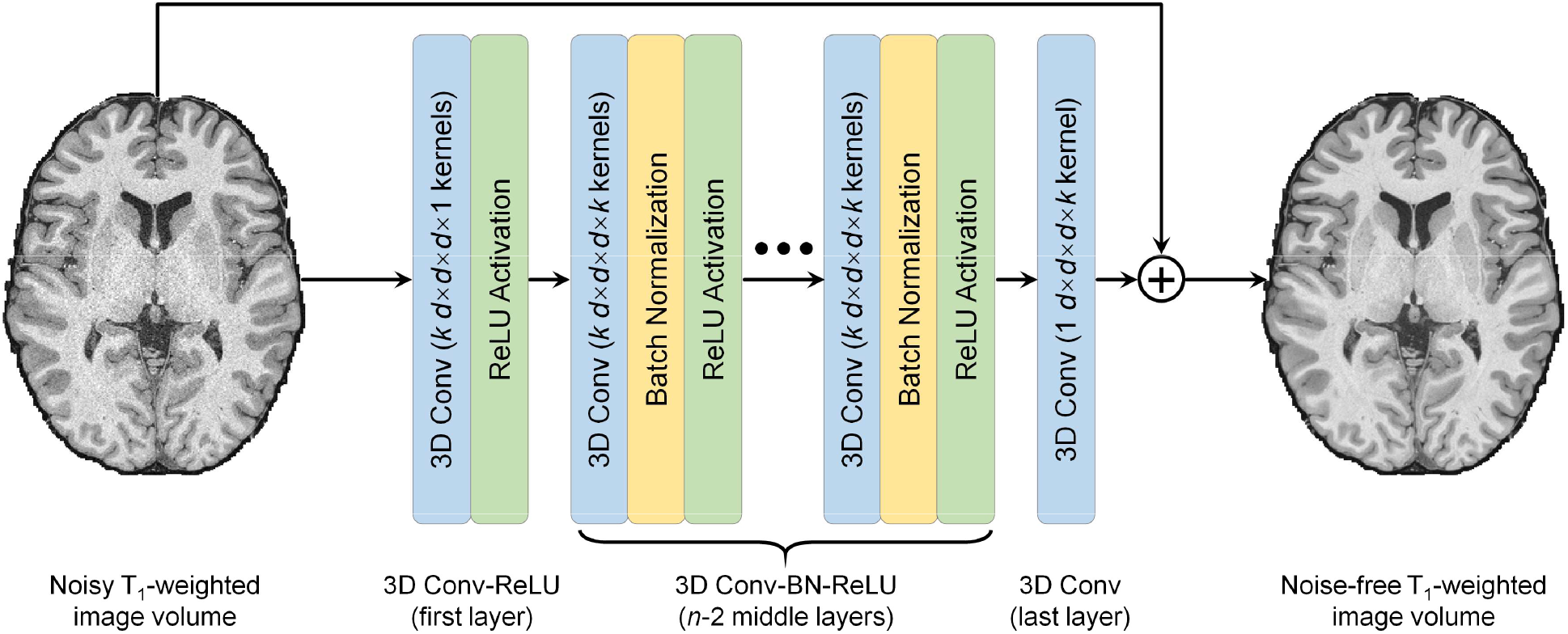
Denoising convolutional neural network (DnCNN) architecture. DnCNN is comprised of stacked 3-dimenstional convolutional filters with rectified linear unit (ReLU) activation functions and batch normalization functions (*n*=20, *k*=64, *d*=3). The input is a noisy anatomical (i.e., T_1_-weighted) image volume from one repetition of the sub-millimeter acquisition. The output is a high-signal-to-noise ratio T_1_-weighted image volume obtained by averaging data from multiple repetitions of the same sub-millimeter acquisition. DnCNN maps the input image volume to the residual volume between the input and output image volume (i.e., noise).

Our DnCNN network was implemented using the Keras application program interface (https://keras.io) with a Tensorflow backend (Google, Mountain View, California, https://www.tensorflow.org). Training and validation were performed on image blocks of 81×81×81 voxels (12 blocks per whole-brain volume) because of the limited GPU memory available. Sixty blocks from five random repetitions of data from each of the four training subjects (240 blocks in total) were used for training, while 12 blocks from the held-out repetition of data from each of 4 training subjects (48 blocks in total) were used for validation. The input and output image volumes were skull-stripped and image intensities were standardized by subtracting the mean intensity and then divided by the standard deviation of the image intensities of the brain voxels from the input noisy single-repetition image volumes.

All 3-dimensional convolutional kernels were randomly initialized with a “He” initialization^110^. Network parameters were optimized by minimizing the mean squared error within the brain (excluding the cerebellum which is irrelevant for cerebral cortical surface reconstruction and has significantly different texture compared to the cerebrum) using stochastic gradient descent^111^ with an Adam optimizer^112^ (default parameters except for the learning rate) using a V100 graphics processing unit (GPU, 16G memory) (NVidia, Santa Clara, California) for about ∼15 hours. The batch size was set to 1, the largest batch the GPU memory can accommodate. The learning rate was 0.0001 for the 20 epochs, after which the network approached convergence, and 0.00005 for another 10 epochs (∼0.5 hour per epoch).

The trained DnCNN was applied to the skull-stripped and standardized whole-brain image volumes from each repetition of the 5 evaluation subjects. The predicted image intensities were transformed to the original range by multiplying with the standard deviation of the image intensities and adding the mean intensity of the brain voxels from the input noisy image volumes.

### Surface reconstruction

Cortical surface reconstruction at native 0.6-mm isotropic resolution was performed using the FreeSurfer sub-millimeter pipeline^43^ (version v6.0) (https://surfer.nmr.mgh.harvard.edu/fswiki/SubmillimeterRecon). The “*recon-all*” function was executed with the “*hires*” option with an expert file that specifies 100 iterations of surface inflation in the “*mris_inflate*” function.

### Image similarity comparison

To quantify the similarity between the image volumes from the single-repetition and 2- to 5-repetition averaged data and the denoised image volumes compared to the high-SNR image volumes from the 6-repetition averaged data, the mean absolute difference (MAD), peak SNR (PSNR) and structural similarity index (SSIM)^113^ were computed using custom scripts written for the MATLAB software package (MathWorks, Natick, Massachusetts). PSNR, defined as 10log_10_(maximum possible value^2^/mean squared error), was computed using MATLAB’s “*psnr*” function, with larger value indicating lower mean squared error. SSIM is a perceptual metric for image similarity quantification ranging between 0 and 1 with larger value indicating higher similarity. SSIM was computed using MATLAB’s “*ssim*” function. For these calculations, the standardized image intensities, which virtually fell into the range [−3, 3], were normalized to the range of [0, 1] by adding 3 and dividing by 6. The group-level mean and standard deviation of the whole-brain averaged MAD, PSNR and SSIM (only within the brain mask) across 30 image volumes from the 5 evaluation subjects were computed to summarize the overall image similarity.

### Image contrast comparison

The gray–white contrast is essential for successful cortical surface reconstruction. It was therefore computed from different types of images and compared. Specifically, the gray–white contrast was computed at the location of every vertex on the gray–white surface reconstructed from the 6-repetition averaged images as [white – gray]/[white + gray]⋅100%. The white matter intensity was sampled at 0.6 mm (i.e., the voxel size) below the gray–white surface. The gray matter intensity was sampled at 0.6 mm above the gray–white surface. The group-level mean and standard deviation of the whole-brain averaged vertex-wise gray–white contrast and variation of the vertex-wise gray–white contrast across 30 image volumes from the 5 evaluation subjects were computed to summarize the overall image contrast level.

### Surface quality comparison

Percentage of defective vertices and surface smoothness were used for assessing the surface quality. The percentage of defective vertices was computed as the number of vertices identified as being part of topological defects in the initial reconstructed surface prior to topology correction^114^ (identified by FreeSurfer surface results files “*lh.defect_labels*” and “*rh.defect_labels*”) divided by the total number of vertices. The surface smoothness was defined as the median value of the whole-brain vertex-wise mean curvature, which was computed using FreeSurfer’s “*mris_curvature*” function. The group-level mean and standard deviation of the percentage of defective vertices, gray–white surface smoothness, gray–CSF surface smoothness across 30 image volumes from the 5 evaluation subjects were computed to summarize the overall surface quality.

### Surface accuracy comparison

The FreeSurfer longitudinal pipeline^101,115,116^ (https://surfer.nmr.mgh.harvard.edu/fswiki/LongitudinalProcessing) was used to quantify the discrepancies of the gray–white and gray–CSF surface positioning and cortical thickness estimation from the single-repetition data, 2-to 5-repetition averaged data, and denoised single-repetition data compared to those from the 6-repetition averaged data. For any two image volumes for comparison, the longitudinal pipeline provides a vertex correspondence between the cortical surfaces reconstructed for each image volume, which enables the calculation of a per-vertex vertex-wise displacement for the two bounding surfaces of the cortical gray matter as well as a per-vertex difference of the cortical thickness estimates^43,117^. The group-level mean and standard deviation of the mean absolute gray–white and gray–CSF surface displacement and cortical thickness difference averaged across the whole brain as well as within 34 cortical parcels (left and right hemispheres combined) from the Desikan-Killiany Atlas provided by FreeSurfer across 30 image volumes from the 5 evaluation subjects were computed to summarize the overall surface positioning accuracy.

### Surface precision comparison

The FreeSurfer longitudinal pipeline was used to quantify the discrepancies of the gray–white and gray–CSF surface positioning and cortical thickness estimation from two consecutively acquired single-repetition data (5 comparisons in total, i.e., repetition 1 vs. repetition 2, repetition 2 vs. repetition 3 to repetition 5 vs. repetition 6) denoised by DnCNN, BM4D and AONLM as described in the previous section to quantify the scan-rescan reproducibility. The group-level mean and standard deviation of the mean absolute gray–white and gray–CSF surface displacement and cortical thickness difference averaged across the whole brain as well as within 34 cortical parcels (left and right hemispheres combined) from the Desikan-Killiany Atlas provided by FreeSurfer across 25 image volumes from the 5 evaluation subjects were computed to summarize the overall surface positioning precision.

### Statistical comparison

The non-parametric Wilcoxon signed-rank test was performed for two related samples to evaluate differences between metrics of interest (i.e., the whole-brain averaged MAD, PSNR, SSIM, vertex-wise gray–white contrast, variability of the vertex-wise gray–white contrast, gray–white surface displacement, gray–CSF surface displacement and cortical thickness difference) from different types of image volume (e.g., DnCNN-denoised data vs. BM4D-denoised data, and DnCNN-denoised data vs. 6-repetition averaged data) across 30 image volumes from the 5 evaluation subjects (*N*=30) for the accuracy metrics and across 25 images volume from the 5 evaluation subjects (*N*=25) for the precision metrics.

## Results

Figure 2 shows that the T_1_-weighted images (Fig. 2, rows a, b, columns iii–v) denoised using DnCNN, BM4D and AONLM were better able to delineate fine anatomical structures compared to the single-repetition data (Fig. 2, rows a, b, columns ii) and were visually more comparable to the T_1_-weighted image generated from the 6-repetetion averaged data (Fig. 2, rows a, b, column i). The textural details were more clearly displayed with sharper gray–white and gray–CSF boundaries in the denoised images, especially in the DnCNN-denoised images, compared to those from the single-repetition data (Fig. 2, row b, arrow heads). The differences between the denoised images and 6-repetition averaged image exhibited negligible levels of anatomical structure and were substantially lower compared to those from the single-repetition images (Fig. 2, row c), with lower whole-brain averaged MAD (0.014, 0.015, 0.015 for DnCNN, BM4D and AONLM vs. 0.03) and higher PSNR (34.7 dB, 34 dB and 34 dB for DnCNN, BM4D and AONLM vs. 28.18 dB). The SSIM maps demonstrate that the denoised images were also perceptually similar to the 6-repetition averaged image (Fig. 2, row d), with higher whole-brain averaged SSIM (0.94, 0.93, 0.93 for DnCNN, BM4D and AONLM vs. 0.81). Overall, DnCNN had slightly better denoising performance than BM4D and AONLM, which generated images of very similar quality to one another.

**Figure 2.**
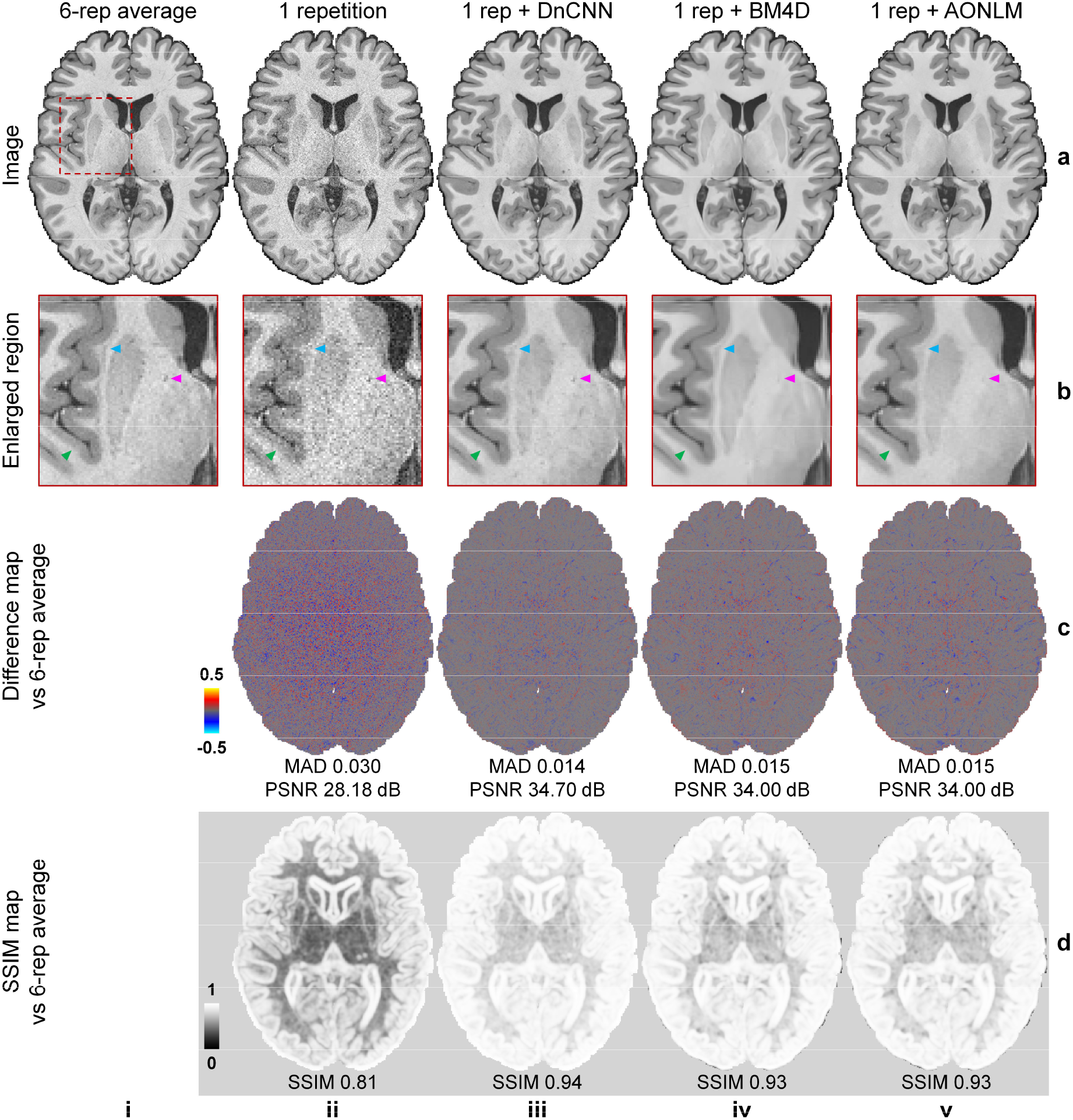
Image quality and similarity. A representative axial image slice from the 6-repetition averaged data (a, i), the single-repetition data (a, ii), and the single-repetition data denoised by DnCNN (a, iii), BM4D (a, iv) and AONLM (a, v), along with magnified views of basal ganglia (row b). The arrowheads highlight the boundary of gray matter and white matter (green), the claustrum (blue), and the blood vessel in the internal capsule (magenta). The difference maps (row c) and the structural similarity index (SSIM) maps (row d) depict the similarity between the images from the single-repetition data and the denoised data compared to the images from the 6-repetition averaged data. SSIM scores range between 0 and 1, with larger values indicating higher similarity. The whole-brain mean absolute difference (MAD), mean peak signal-to-noise ratio (PSNR) and mean SSIM are used to quantify the image similarity.

The noise estimated by the three methods (i.e., the residual maps between the denoised images and the acquired single-repetition image) did not contain any noticeable anatomical structure or biases reflecting the underlying anatomy (Supplementary Fig. 2, rows b, columns iii-v) and were visually similar to the residual map from the 6-repetition averaged data (Supplementary Fig. 2, rows b, column ii). DnCNN estimated slightly lower noise levels in the center of the brain compared to BM4D and AONLM.

The similarity between images from the single-repetition data and the denoised single-repetition data compared to the 6-repetition averaged data was quantified at the group level and compared to those from the 2- to 5-repetition averaged data in Figure 3. The group-level means (± the group-level standard deviation) of the whole-brain averaged MAD, PSNR and SSIM of the denoised images were approximately two times lower (0.0153 ± 0.0018, 0.0165 ± 0.0017, 0.0163 ± 0.0017 for DnCNN, BM4D and AONLM vs. 0.0329 ± 0.0035), ∼6 dB higher (34.0 ± 0.98 dB, 33.3 ± 0.83 dB, 33.4 ± 0.85 dB for DnCNN, BM4D and AONLM vs. 27.5 ± 0.89 dB) and approximately 0.12 higher (0.93 ± 0.012, 0.92 ± 0.011, 0.92 ± 0.011 for DnCNN, BM4D and AONLM vs. 0.80 ± 0.028), respectively, compared to the single-repetition images. The image similarity levels of the denoised images compared to the 6-repetition averaged images were equivalent to those derived from what would be expected from averaging 2.5 images (2.72, 2.53, 2.55 in terms of MAD, 2.63, 2.42, 2.44 in terms of PSNR, and 2.55, 2.22, 2.23 in terms of SSIM for DnCNN, BM4D and AONLM).

**Figure 3.**
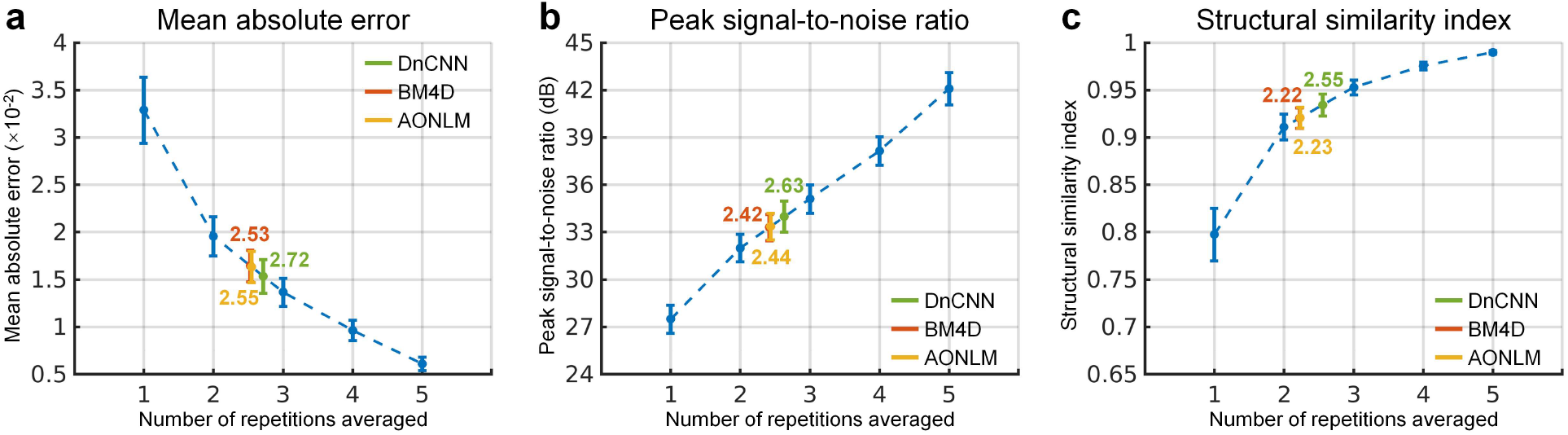
Image similarity quantification. The blue dots and error bars represent the group mean and standard deviation of the mean absolute difference (MAD) (a), peak signal-to-noise ratio (PSNR) (b) and structural similarity index (SSIM) (c) of the images from the single-repetition data and 2- to 5-repetition averaged data compared to the images from the 6-repetition averaged data across 30 image volumes from the 5 evaluation subjects. The green, red and yellow dots and error bars represent the group mean and standard deviation of the MAD, PSNR and SSIM of the images from the single-repetition data denoised by DnCNN (green), BM4D (red) and AONLM (yellow) compared to the images from the 6-repetition averaged data across 30 image volumes from the 5 evaluation subjects. The red dots and error bars for BM4D are virtually covered by the yellow dots and error bars for AONLM. The numbers above and below the dashed lines that link two neighboring blue dots indicate the number of image volumes that would be needed to obtain equivalent similarity metrics to those obtained from images from single-repetition data denoised by different methods. All comparisons of results from different denoising methods and numbers of repetitions for averaging are statistically significant (*p*<0.05).

The vertex-wise gray–white contrast maps depict the spatial variation of the gray–white contrast across cortical regions (Fig. 4, averaged maps across 30 image volumes from the 5 evaluation subjects available in Supplementary Fig. 3). Overall, the gray–white contrast was consistently lower in heavily myelinated regions near the central sulcus, calcarine sulcus, paracentral gyrus, Heschl’s gyrus, posterior cingulate gyrus, isthmus of the cingulate gyrus with increased gray matter image intensity on T_1_-weighted images, and in the insula with decreased white matter image intensity on the T_1_-weighted images arising from the partial volume effects with the claustrum^47^. These regions are more prone to noise during the cortical surface reconstruction process.

**Figure 4.**
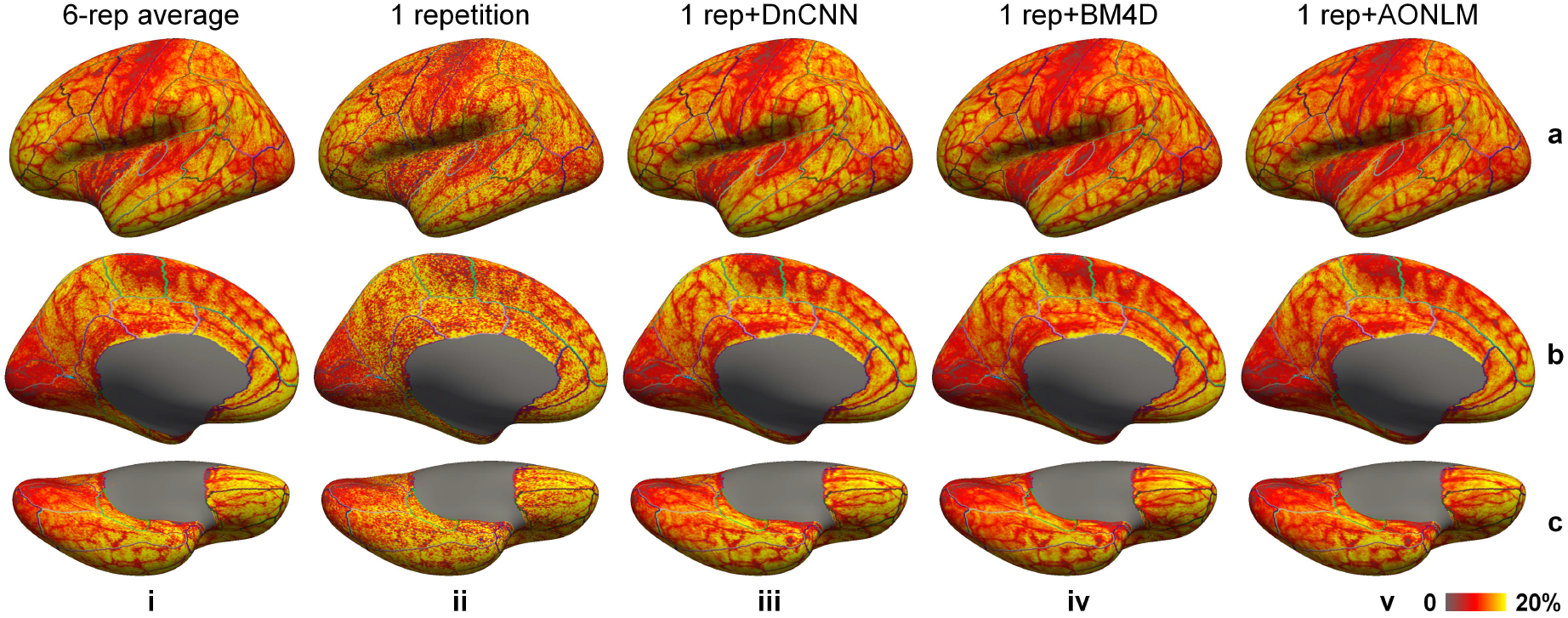
Gray–white contrast. Left-hemispheric vertex-wise contrast between the gray matter and white matter image intensity (defined as [white − gray]/[white + gray]⋅100%) from the 6-repetition averaged data (column i), single-repetition data (column ii), single-repetition data denoised by DnCNN (column iii), BM4D (column iv) and AONLM (column v) from a representative subject, displayed on inflated surface representations. Different cortical regions from the FreeSurfer cortical parcellation (i.e., *aparc.annot*) are depicted as colored outlines

Gray–white contrast is critical for accurate cortical surface reconstruction, which should be preserved as much as possible during the noise removal process. The gray–white contrast map from the single-repetition data (Fig. 4, column ii) displayed similar gray–white contrast level compared to the map from the 6-repetition averaged data (Fig. 4, column i), but was more spatially variable. The gray–white contrast maps from the denoised data (Fig. 4, columns iii–v) were smoother and less spatially variable, appearing more similar to the map from the 6-repetition averaged data (Fig. 4, column i). The gray–white contrast level in the map from DnCNN (Fig. 4, column iii) was slightly higher than those from BM4D (Fig. 4, column iv) and AONLM (Fig. 4, column v), and visibly more similar to that from the 6-repetition averaged data (Fig. 4, column i).

The group-level means (± the group-level standard deviation) of the spatial mean and spatial variability of the vertex-wise gray–white contrast across the whole brain were quantified to compare the capability of different denoising methods in preserving gray–white contrast (Fig. 5). The whole-brain averaged vertex-wise gray–white contrast stayed almost the same and decreased only marginally when averaging more repetitions of data (Fig. 5a, blue curve), which was consistent with visual inspection (Fig. 4, columns i, ii). The whole-brain averaged vertex-wise gray–white contrast from DnCNN was slightly higher than that from BM4D and AONLM (13.03 ± 0.71 vs. 12.20 ± 0.73 and 12.10 ± 0.76 for BM4D and AONLM) and was only slightly lower than that from the 6-repetition averaged data (13.03 ± 0.71 vs. 13.30 ± 0.59).

**Figure 5.**
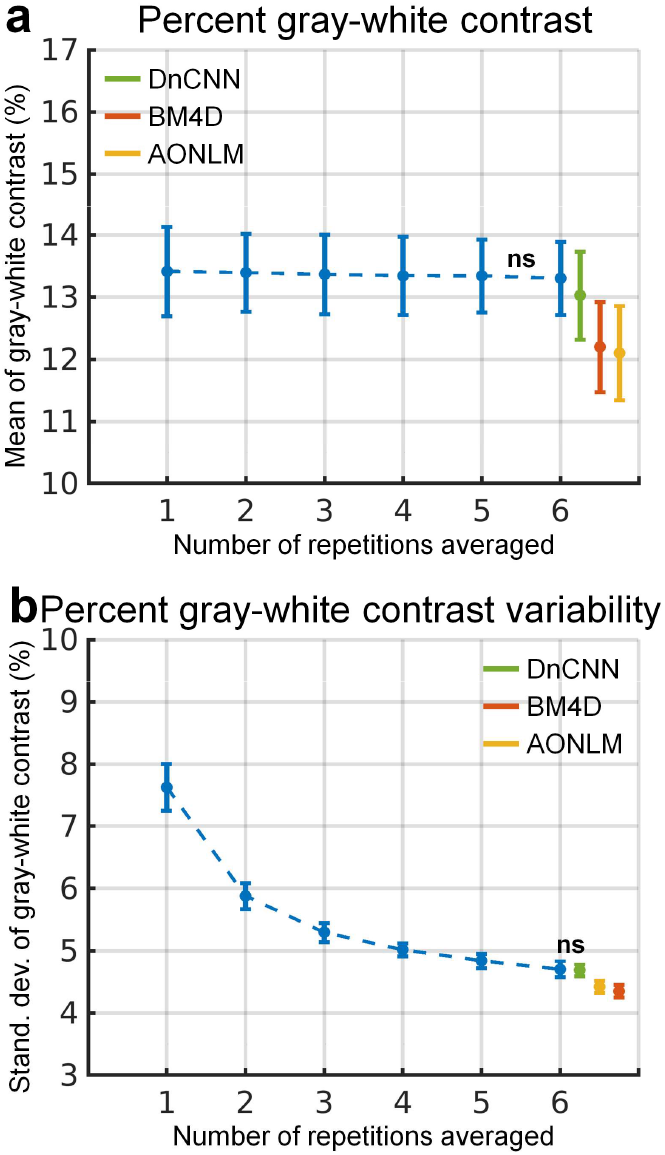
Gray–white contrast quantification. The blue dots and error bars represent the group mean and standard deviation of the spatial mean of the vertex-wise percent gray–white contrast (defined as [white - gray]/[white + gray]⋅100%) across the whole-brain cortical surface (a) and the spatial variability (calculated as standard deviation) of the vertex-wise percent gray–white contrast across the whole-brain cortical surface (b) of the images from the single-repetition data and 2- to 6-repetition averaged data across 30 image volumes from the 5 evaluation subjects. The green, red and yellow dots and error bars represent the group mean and standard deviation of the spatial mean value and spatial variability of the percent gray-white contrast of the images from the single-repetition data denoised by DnCNN (green), BM4D (red) and AONLM (yellow) across 30 image volumes from the 5 evaluation subjects. All comparisons of results from different denoising methods and numbers of repetitions for averaging are statistically significant (*p*<0.05), except for between the 5- and 6-repetition averaged data for the spatial mean value of percent gray–white contrast (a), and between the 6-repetition averaged data and the single-repetition data denoised by DnCNN for the spatial variability of the percent gray-white contrast (b) (denoted with abbreviation “ns”).

The spatial variability of the vertex-wise gray–white contrast continued to decrease when more repetitions of the data were averaged (Fig. 5b, blue curve). This finding was also visually reflected by smoother gray–white contrast maps (Fig. 4, columns i, ii). The spatial variability of the vertex-wise gray–white contrast in the DnCNN-denoised data was nearly the same as that in the 6-repetition averaged data (4.69 ± 0.10 vs. 4.70 ± 0.13, comparison not significant). The spatial variability in the BM4D- and AONLM-denoised data (4.35 ± 0.10 vs. 4.42 ± 0.10) was further reduced than in the DnCNN-denoised data and 6-repetition averaged data, potentially due to stronger unwanted spatial smoothing effects that reduce tissue contrast.

The effects of data averaging and denoising on the reconstructed cortical surfaces are shown in Figure 6. The cortical surfaces from the single-repetition data (Fig. 6, column ii) resembled the overall cortical geometry represented by the surfaces from the 6-repetition averaged data (Fig. 6, column i), but were not smooth. The bumpy appearance was more obviously displayed in the superior and inferior brain (Fig. 6, column ii, arrow heads), where the SNR was further reduced because of the lower signal level resulting from the non-uniform slab excitation. The cortical surfaces from the 6-repetition averaged data and the denoised data were much smoother and more visually similar (Fig. 6, column i, iii–v). The gray–CSF surfaces were in general smoother compared to the gray–white surfaces.

**Figure 6.**
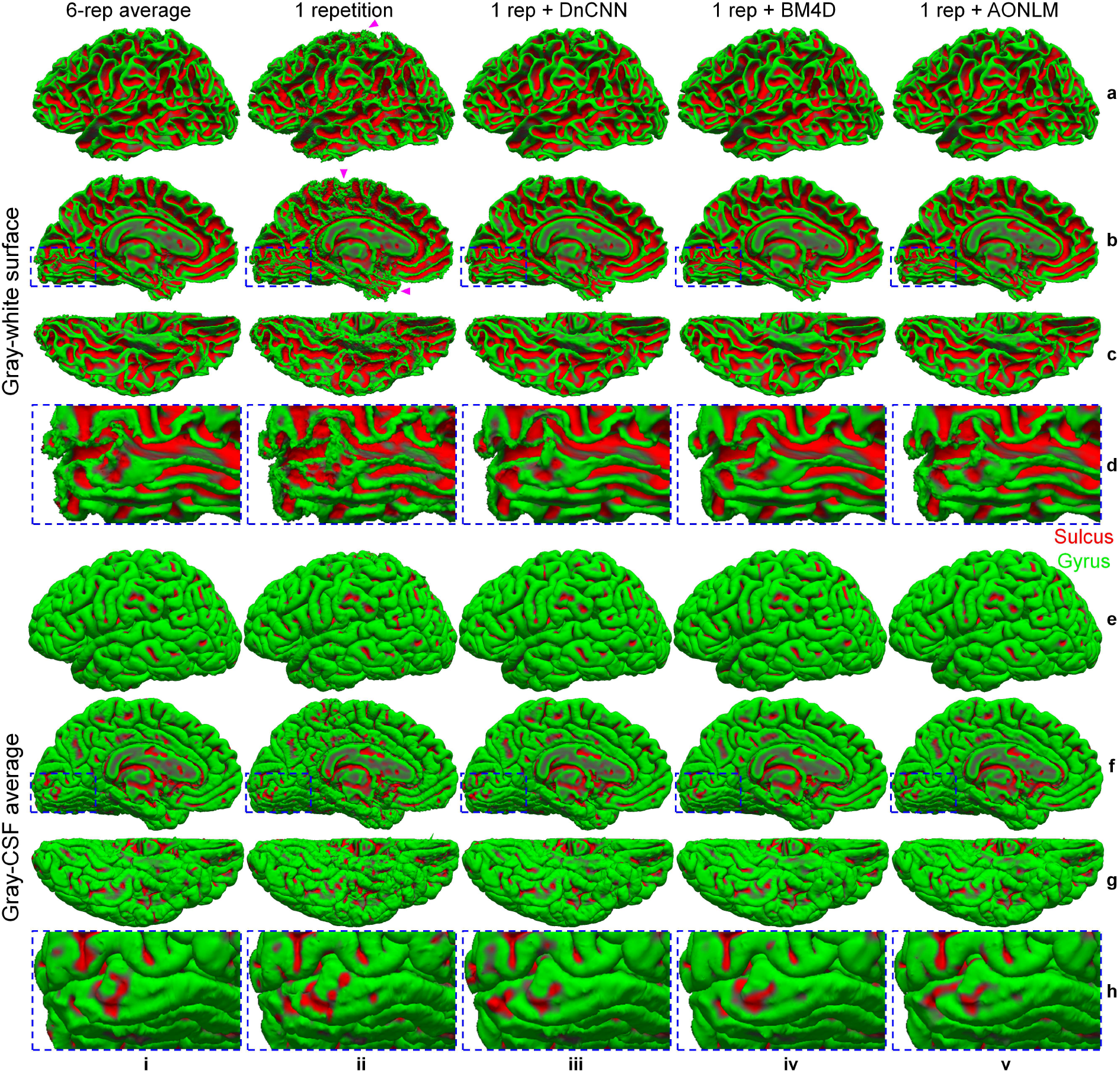
Cortical interface surface. Gray–white interface surfaces (a–d) and gray–CSF interface surfaces (e–h), along with magnified views near the Calcarine sulcus (rows d, h), reconstructed from the 6-repetition averaged data (column i), the single-repetition data (column ii), and the single-repetition data denoised by DnCNN (column iii), BM4D (column iv) and AONLM (column v) from the left hemisphere of a representative subject. The arrow heads highlight locations toward the superior and inferior brain with reduced signal-to-noise ratio in the acquired data because of the non-uniform slab excitation profile and consequently less smooth cortical surfaces.

The mean curvature can be used to characterize the local smoothness of cortical surfaces quantitatively (Fig. 7, averaged maps across 30 image volumes from the 5 evaluation subjects available in Supplementary Fig. 4)—the mean curvature map of a smooth surface will exhibit low levels of spatial variability. For the single-repetition data, the mean curvature map of the gray–white surface was highly spatially variable near the central sulcus, calcarine sulcus, paracentral gyrus, cingulate gyrus and insula, where the gray–white contrast was lower (Fig. 4) and the image noise had a higher impact on the reconstructed cortical surfaces. The mean curvature maps from the 6-repetition averaged data and the denoised data were much less spatially variable, but were still slightly variable near the superior portion of the central sulcus because of the simultaneous reduction in gray–white contrast and lower signal levels near the edge of the excited slab.

**Figure 7.**
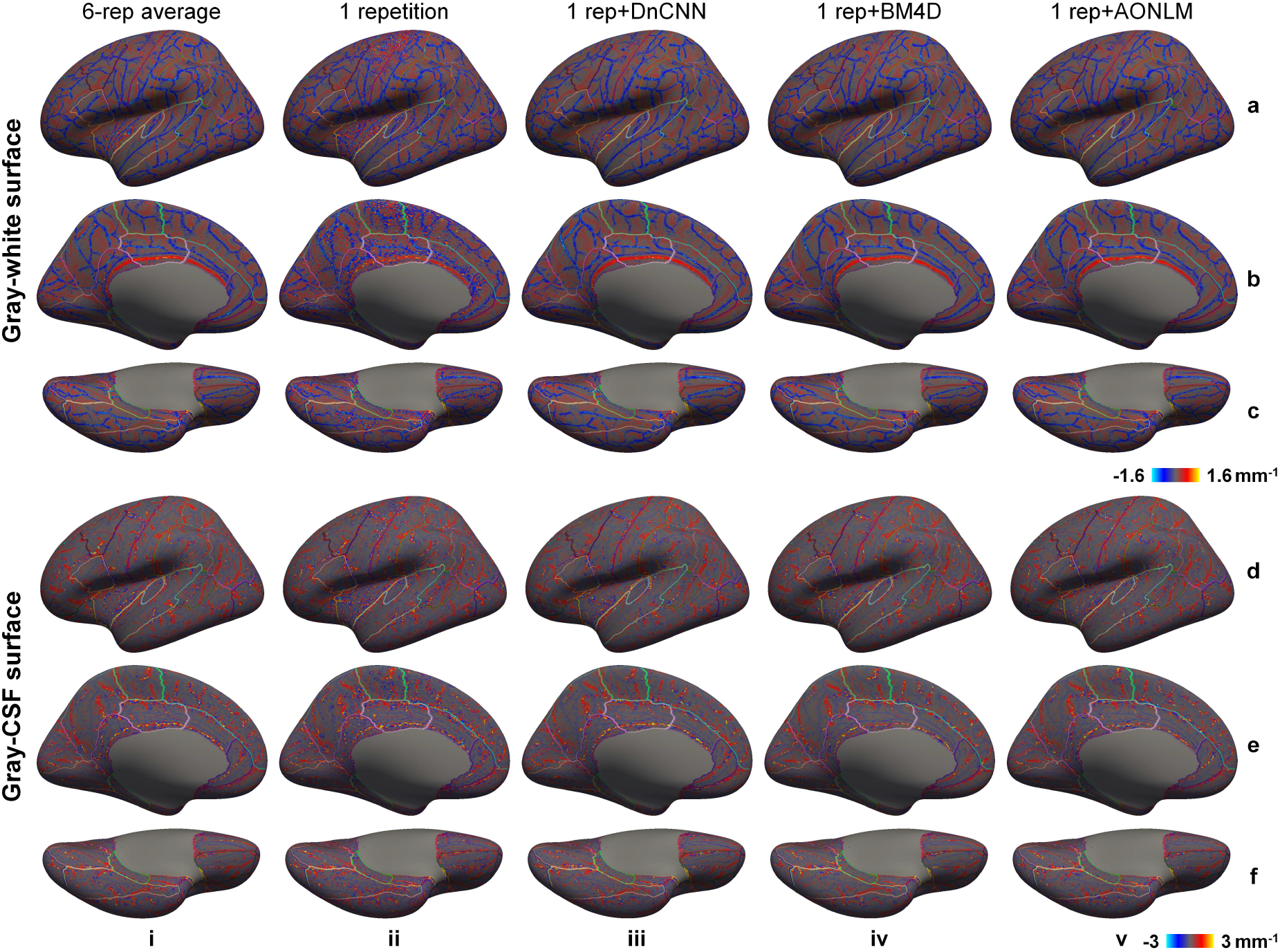
Cortical surface smoothness. Left-hemispheric vertex-wise mean curvature of the reconstructed gray–white surfaces (a–c) and gray–CSF surfaces (d–f) from the 6-repetition averaged data (column i), single-repetition data (column ii), single-repetition data denoised by DnCNN (column iii), BM4D (column iv) and AONLM (column v) from a representative subject, displayed on inflated surface representations. Different cortical regions from the FreeSurfer cortical parcellation (i.e., *aparc.annot*) are depicted as colored outlines

Quantitatively, the median of the whole-brain mean curvature decreased as the number of the averaged repetition increased (Fig. 8a, b, blue curves), indicating increased overall surface smoothness. The cortical surfaces from the denoised data were even smoother than those from the 6-repetition data (0.17 ± 0.0051, 0.16 ± 0.0042 and 0.16 ± 0.0052 for DnCNN, BM4D and AONLM vs. 0.18 ± 0.0061 for 6-repetition averaged data for gray–white surface; 0.19 ± 0.0039, 0.18 ± 0.0039 and 0.18 ± 0.0035 for DnCNN, BM4D and AONLM vs. 0.20 ± 0.0038 for 6-repetition averaged data for gray–white surface), potentially due to stronger unwanted spatial smoothing effects. The surface smoothness from DnCNN was more similar to the surface smoothness from the 6-repetition averaged data compared to those from BM4D and AONLM. The percentage of vertices identified as being part of topological defects in the initial surfaces reconstructed from the input data prior to automatic topological correction decreased as the number of the averaged repetitions increased (Fig. 8c, blue curve), indicating improved image quality due to data averaging. The percentage of the defective vertices (typically 0.8 to 1 million vertices in total for a subject for 0.6-mm isotropic data) was even lower in the denoised data than in the 6-repetition averaged data (0.014% ± 0.0028%, 0.01% ± 0.0015% and 0.01% ± 0.0017% for DnCNN, BM4D and AONLM vs. 0.023% ± 0.0056% for 6-repetition averaged data).

**Figure 8.**
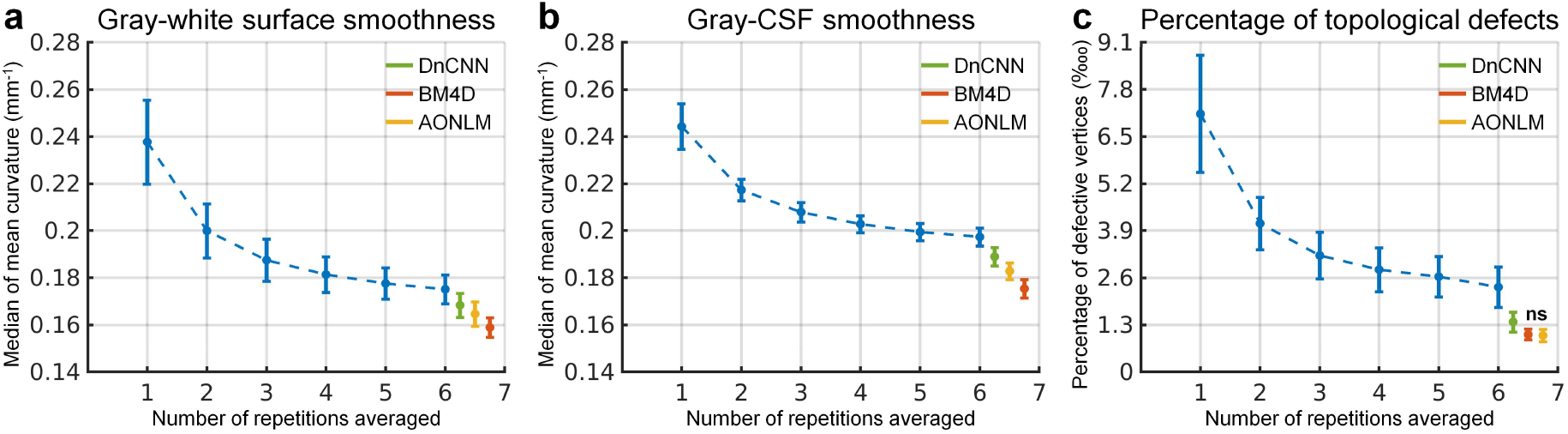
Cortical surface quality quantification. The blue dots and error bars represent the group mean and standard deviation of the smoothness of the gray–white (a) and gray–CSF surface (b) (defined as the median of the vertex-wise mean curvature across the cortical surface), and the percentage of the vertices identified as being part of topological defects in the initial surface (c) from the single-repetition data and 2- to 6-repetition averaged data across 30 image volumes from the 5 evaluation subjects. The green, red and yellow dots and error bars represent the group mean and standard deviation of the gray–white and gray–CSF surface smoothness and percentage of the defective vertices from the single-repetition data denoised by DnCNN (green), BM4D (red) and AONLM (yellow) across 30 image volumes from the 5 evaluation subjects. All comparisons of results from different denoising methods and numbers of repetitions for averaging are statistically significant (p<0.05), except for between the BM4D- and AONLM-denoised data for the percentage of the defective vertices (c) (denoted with abbreviation “ns”).

To demonstrate the effects of data averaging and denoising on the accuracy of surface positioning, cross-sections of the cortical surfaces were presented in enlarged views near the central sulcus and the calcarine sulcus (Fig. 9), chosen because they are regions with reduced gray–white contrast. The cortical surfaces reconstructed from the denoised data were similar to those reconstructed from 6-repeition averaged data and were markedly improved compared to the surface placement derived from the single-repetition data (Fig. 9, arrow heads), especially the gray–white surfaces. The discrepancy in positioning of the gray–CSF surface across the different denoising techniques was overall smaller than that of the gray–white surface.

**Figure 9.**
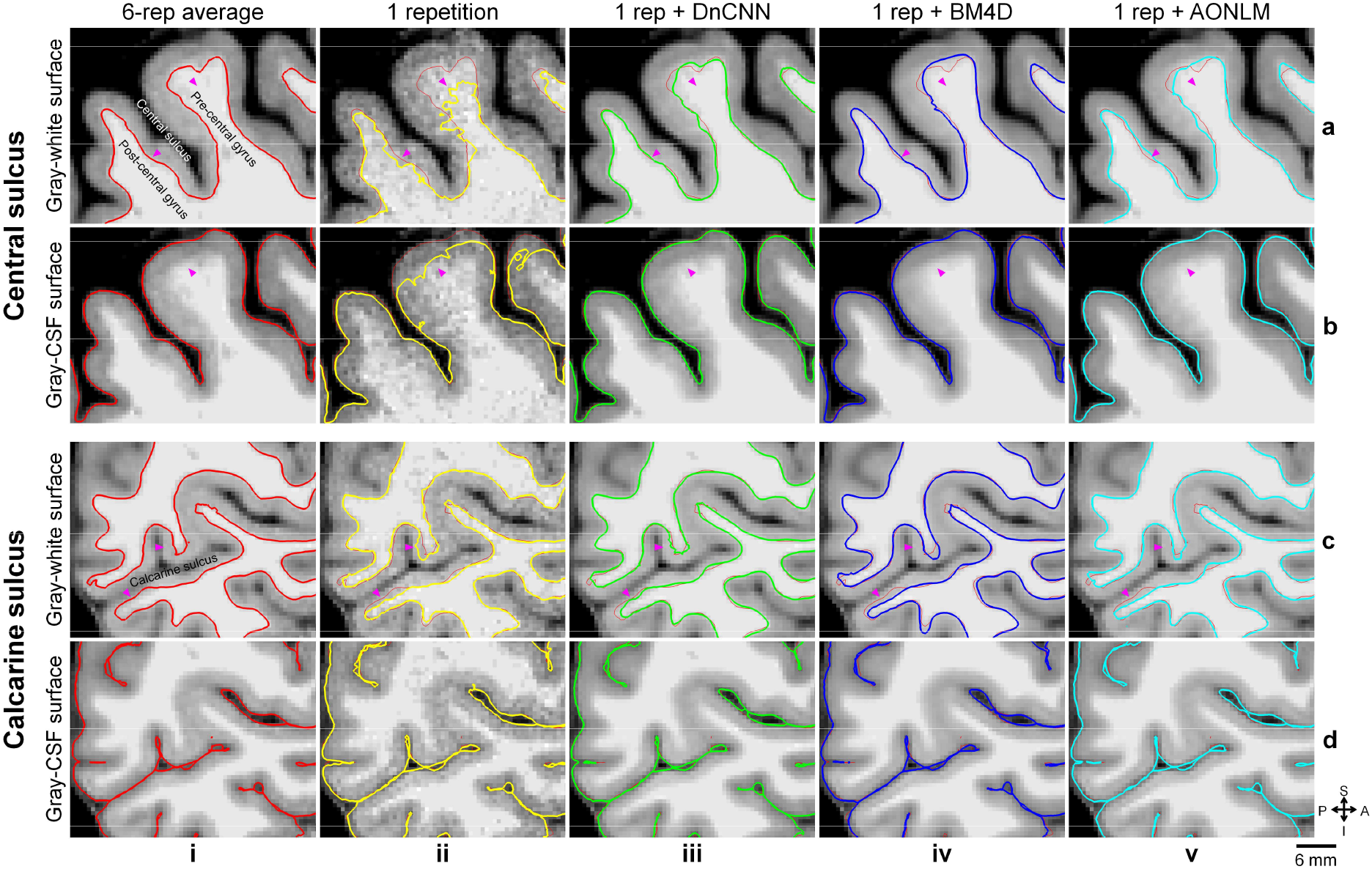
Cortical surface cross-sections. Enlarged views of sagittal image slices from the 6-repetition averaged data (column i), the single-repetition data (column ii), and the single-repetition data denoised by DnCNN (column iii), BM4D (column iv) and AONLM (column v) near the central sulcus (rows a, b) and the calcarine sulcus (rows c, d) from the left hemisphere of a representative subject. Cross-sections of the gray–white (rows a, c) and gray–CSF surfaces (rows b, d) are visualized as colored contours and overlaid on top of the images used for reconstructing the respective surfaces. The surface cross-sections from the 6-repetition averaged data are displayed as red thin contours as references along with the surface cross-sections from the single-repetition data and the denoised data (columns ii-v). The arrow heads highlight locations where denoising provides clear improvement in the cortical surface positioning estimates.

The denoised images exhibited substantially lower discrepancy in the surface positioning relative to surfaces reconstructed from the 6-repetition averaged images compared to the discrepancy seen from the single-repetition images in all cortical parcels on the single-subject level (Fig. 10, columns i–iii) and group level (Fig. 10, columns iv–vi). In the denoised results, the discrepancy was most prominent near the central sulcus, calcarine sulcus, Heschl’s gyrus and cingulate gyrus and insula for the gray–white surfaces (Fig. 10, rows a–d, column iv–vi), near the Heschl’s gyrus and insula for the gray–CSF surfaces (Fig. 10, rows e–h, column iv–vi), and near the Heschl’s gyrus and insula for the cortical thickness estimation (Fig. 10, rows i–l, column iv–vi). Estimates from the single-repetition data and denoised data near the temporal pole and orbitofrontal cortex had larger discrepancies presumably because of the large and varying geometric distortions near air-tissue interfaces in each repetition of the data, which could not be accurately aligned for averaging the 6 repetitions of the data and therefore induced increased misalignment between each single-repetition data and the 6-repetition averaged data.

**Figure 10.**
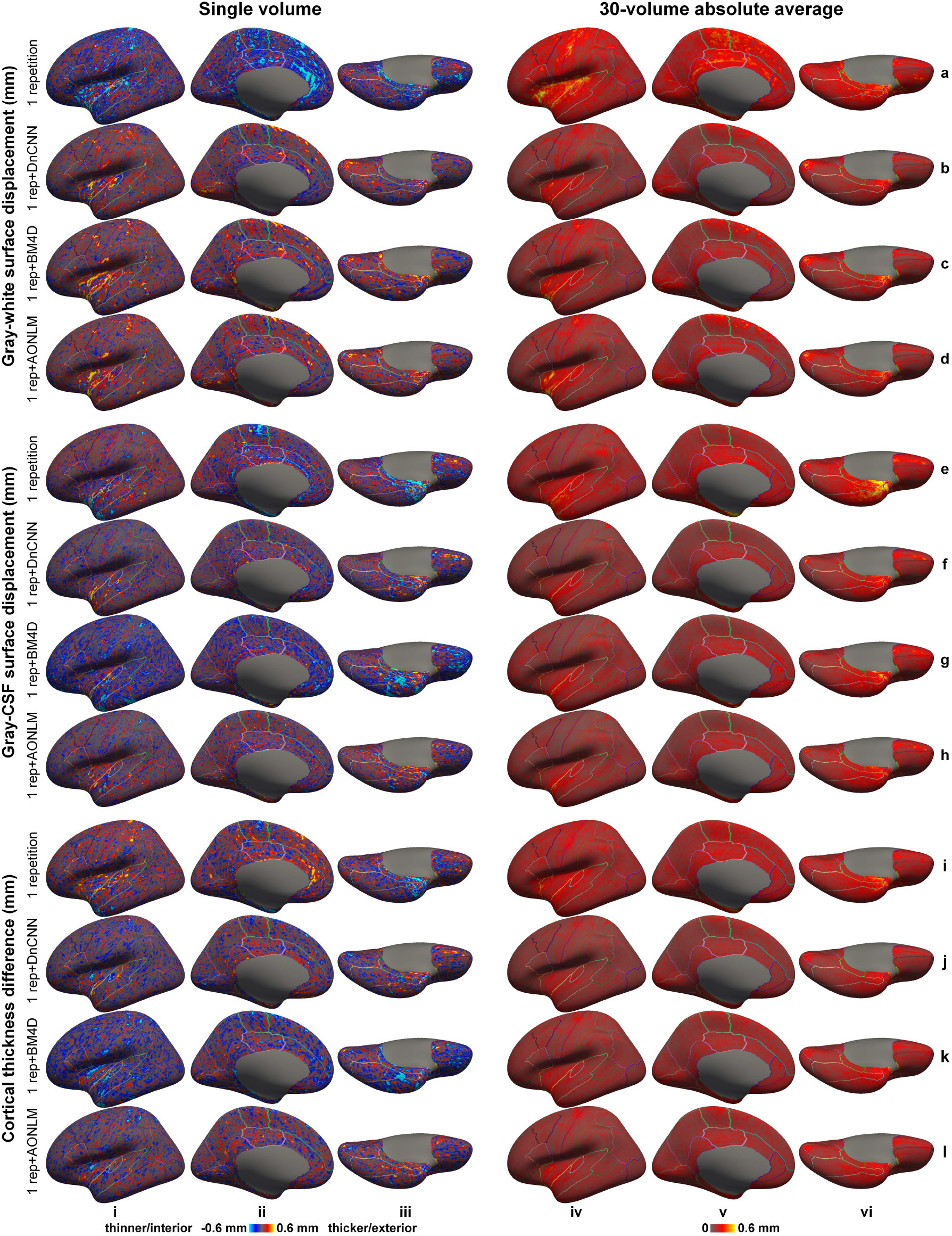
Positioning and thickness accuracy. Left-hemispheric vertex-wise displacement/difference of the gray–white surfaces (rows a–d) and gray–CSF surfaces (rows e–h) and cortical thickness estimates (rows i–l) from the single-repetition data (rows a, e, i) and the single-repetition data denoised by DnCNN (rows b, f, j), BM4D (rows c, g, k) and AONLM (rows d, h, l) compared to the surfaces estimated from the 6-repetition averaged data of a representative subject (column i–iii) and the absolute average across 30 image volumes from the 5 evaluation subjects (rows iv–vi), displayed on inflated surface representations. Different cortical regions from the FreeSurfer cortical parcellation (i.e., *aparc.annot*) are depicted as colored outlines.

The group mean and standard deviation of the mean absolute displacement/difference of the gray–white surface, gray–CSF surface and cortical thickness was quantified for the whole brain (Fig. 11) and each cortical region (Supplementary Tables 1–3). The accuracy of surface positioning and cortical thickness estimation, especially the gray–white surface positioning was increased in the denoised data. The group-level means (± the group-level standard deviation) of the whole-brain averaged displacement of the gray–white surface in the denoised data were about two thirds of those from single-repetition images (0.15 ± 0.02 mm, 0.16 ± 0.025 mm, 0.17 ± 0.025 mm for DnCNN, BM4D and AONLM vs. 0.25 ± 0.04 mm), which were equivalent to those derived from ∼two-repetition averaged images (2.24, 1.92, 1.92 for DnCNN, BM4D and AONLM, respectively). The group-level means (± the group-level standard deviation) of the whole-brain averaged displacement/difference of the gray–CSF surface and cortical thickness in the denoised data were about four fifths of those from single-repetition images (0.13 ± 0.023 mm, 0.15 ± 0.032 mm, 0.14 ± 0.026 mm for DnCNN, BM4D and AONLM vs. 0.19 ± 0.033 mm for gray–white surface positioning; 0.13 ± 0.02 mm, 0.14 ± 0.027 mm, 0.14 ± 0.023 mm for DnCNN, BM4D and AONLM vs. 0.18 ± 0.027 mm for cortical thickness estimation), which were equivalent to those derived from images by averaging 1.5 to 1.9 repetitions of data (1.91, 1.63, 1.78 for DnCNN, BM4D and AONLM for gray–white surface; 1.85, 1.55, 1.64 for DnCNN, BM4D and AONLM for cortical thickness estimation). The improvement for the gray–white surface positioning was slightly larger than for the gray–CSF surface positioning and cortical thickness estimation.

**Figure 11.**
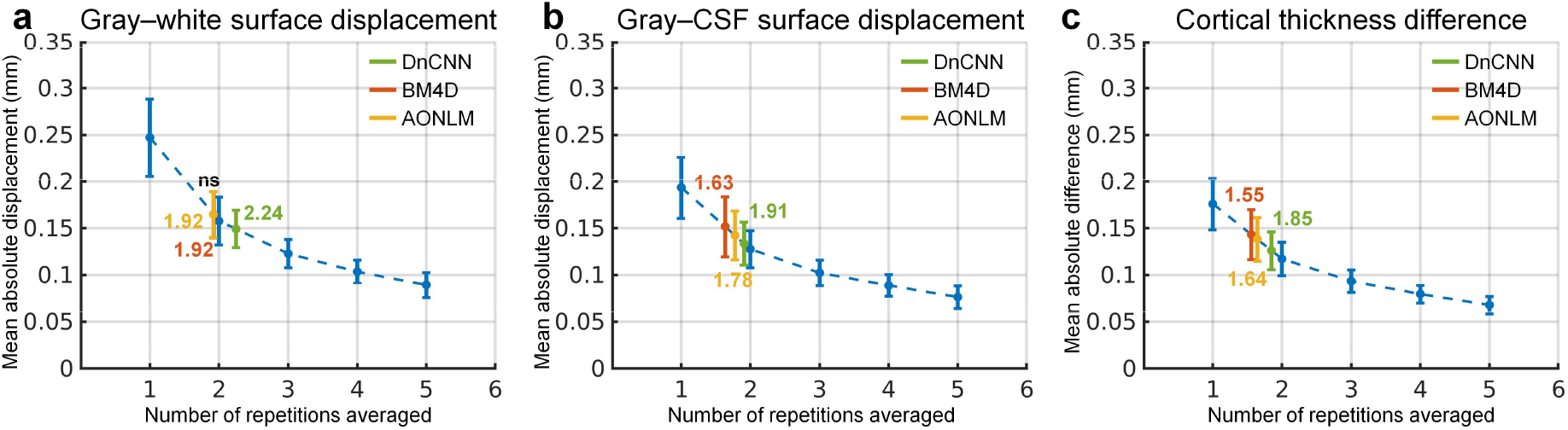
Positioning and thickness accuracy quantification. The blue dots and error bars represent the group mean and standard deviation of the whole-brain averaged absolute displacement/difference of the gray–white surface (a), gray–CSF surface (b) and cortical thickness (c) estimated from the images from the single-repetition data and 2- to 5-repetition averaged data compared to the images from the 6-repetition averaged data across 30 image volumes from the 5 evaluation subjects. The green, red and yellow dots and error bars represent the group mean and standard deviation of the whole-brain averaged absolute displacement/difference for the images from the single-repetition data denoised by DnCNN (green), BM4D (red) and AONLM (yellow) compared to the images from the 6-repetition averaged data across 30 image volumes from the 5 evaluation subjects. The red dots and error bars for BM4D are covered by the yellow dots and error bars for AONLM for the gray–white surface displacement (a). The numbers above and below the dashed lines that link two neighboring blue dots indicate the number of image volumes that would be needed to obtain equivalent similarity metrics to those obtained from images from single-repetition data denoised by different methods. All comparisons of results from different denoising methods and numbers of repetitions for averaging are statistically significant (p<0.05), except for between the BM4D- and AONLM-denoised data for the gray–white surface displacement (a) (denoted with abbreviation “ns”).

The scan-rescan precision of the cortical surface position and thickness estimation using the denoised data was quantified for the whole brain and each cortical region (Fig. 12, Supplementary Table 4). The reproducibility was in general lower near the temporal pole and orbitofrontal cortex presumably because of the elevated image distortions in these regions. The reproducibility was lower near the central sulcus, calcarine sulcus, Heschl’s gyrus and insula for the gray–white surfaces (Fig. 12, rows a–c), near the insula for the gray–CSF surfaces (Fig. 12, rows d–f) and cortical thickness estimation (Fig. 12, rows g–i), regions where accurate surface reconstruction is challenging.

**Figure 12.**
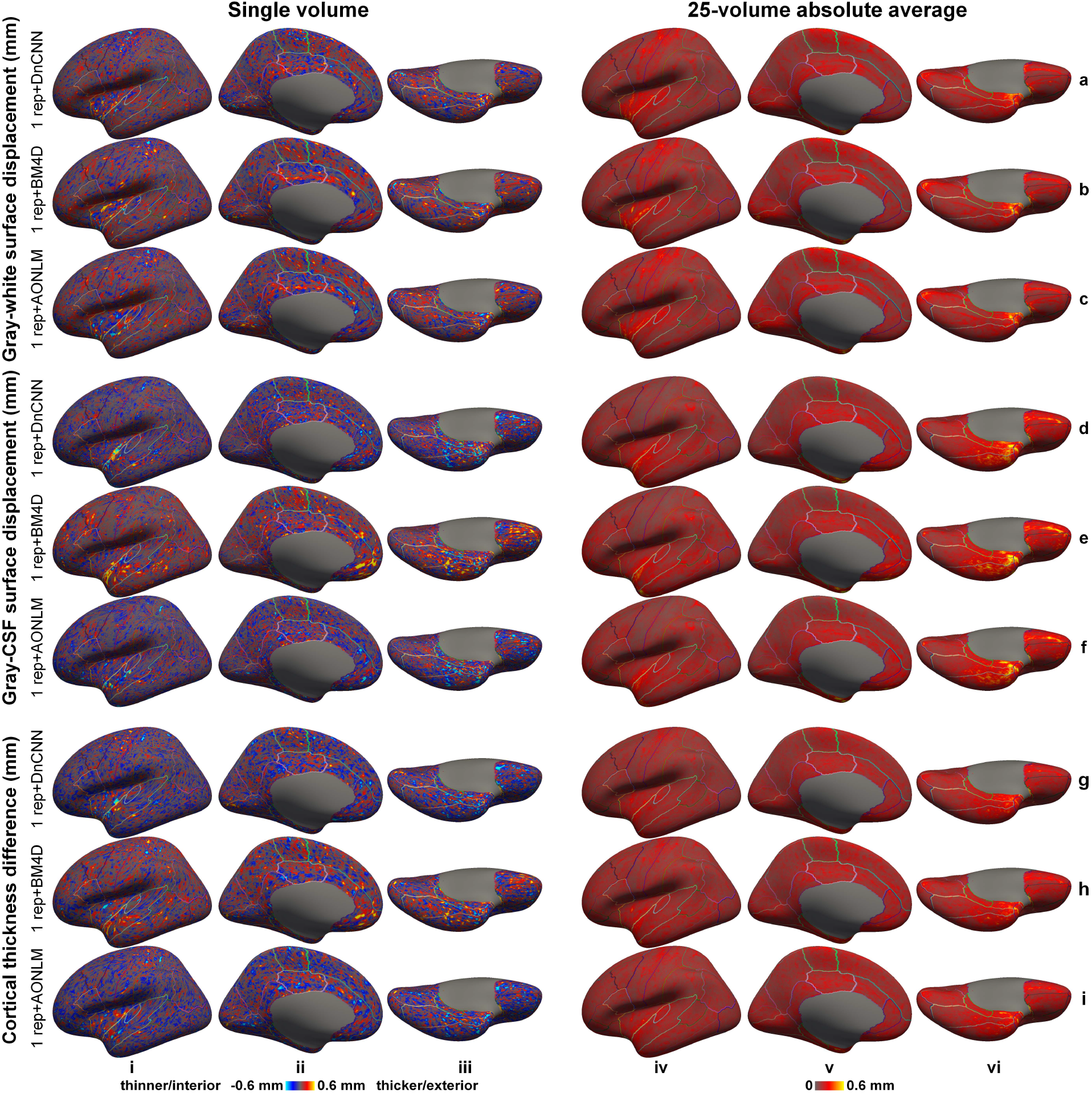
Positioning and thickness precision. Left-hemispheric vertex-wise displacement/difference of the gray–white surfaces (rows a–c) and gray–CSF surfaces (rows d–f) and cortical thickness estimates (rows g–i) from two consecutively acquired single-repetition data denoised by DnCNN (rows a, d, g), BM4D (rows b, e, h) and AONLM (rows c, f, i) of a representative subject (column i–iii) and the absolute average across 25 image volumes from the 5 evaluation subjects (rows iv–vi), displayed on inflated surface representations. Different cortical regions from the FreeSurfer cortical parcellation (i.e., *aparc.annot*) are depicted as colored outlines.

The cortical surface positioning and thickness estimation derived from DnCNN-denoised results were slightly more reproducible than those from BM4D and AONLM. The group mean and standard deviation of the whole-brain averaged absolute displacement/difference between the surface reconstruction results derived from two consecutively acquired single-repetition data denoised by DnCNN, BM4D and AONLM were 0.16 ± 0.034 mm, 0.17 ± 0.034 mm, 0.17 ± 0.035 mm for the gray–white surface, 0.16 ± 0.043 mm, 0.17 ± 0.045 mm, 0.17 ± 0.043 mm for the gray–CSF surface, and 0.15 ± 0.032 mm, 0.16 ± 0.034 mm, 0.16 ± 0.034 mm for the cortical thickness.

## Discussion

In this study, we demonstrate improved *in vivo* human cerebral cortical surface reconstruction at sub-millimeter isotropic resolution using denoised T_1_-weighted images. We show that three well-known classical denoising methods, including DnCNN, BM4D and AONLM, effectively remove the noise in the empirical T_1_-weighted data acquired at 0.6 mm isotropic resolution (∼10 minutes scan) and generate high-SNR T_1_-weighted images similar to those obtained by averaging 6 repetitions (∼ 1 hour scan) of the data with only minimal loss of gray–white image intensity contrast (2% to 9%). The superior quality of the denoised images is reflected by a low whole-brain averaged MAD of ∼0.016, high whole-brain averaged PSNR of ∼33.5 dB and SSIM of 0.92, which is equivalent to those of T_1_-weighted images obtained by averaging ∼2.5 repetitions of the data, saving ∼15 minutes scan time. Due to the significantly reduced noise, the reconstructed cortical surfaces are smoother with similar positioning compared to those from the 6-repetition averaged data. The whole-brain mean absolute difference in the gray–white surface placement, gray–CSF surface placement and cortical thickness estimation is lower than 160 μm, which is equivalent to those of T_1_-weighted images obtained by averaging ∼1.55 to ∼2.24 repetitions of the data, saving ∼5 to ∼12 minutes scan time, and on the same order as the test-retest precision of the FreeSurfer reconstruction (0.1 mm to 0.2 mm)^117–119^.

Our study systematically characterized the effects of denoising on cortical surface reconstruction using empirical data. Three well-known classical denoising methods were evaluated in our study using this characterization framework, similar to what we have applied previously to characterize anatomical image quality in terms of cortical reconstruction precision and accuracy^43,47,117,119^, which can be applied to evaluate other image-based and k-space-based denoising methods and more sophisticated denoising CNNs. We systematically quantified and compared not only the image similarity compared to the ground truth as in most denoising studies, but also the gray–white tissue contrast that is critical for accurate cortical surface reconstruction, surface smoothness and surface placement accuracy from the denoised data. Because our ground-truth data were obtained by data averaging (6 repetitions, ∼1 hour scan), we were able to evaluate the image and surface metrics from the denoised data relatively to those from 2- to 5-repetition averaged data and determine the *efficiency factor*, i.e., the number of image volumes for averaging that would be needed to obtain equivalent image and surface metric values compared to those obtained from denoised single-repetition images. This unique metric concisely and intuitively summarizes the denoising performance of different methods compared to data averaging, the most common practice to increase the SNR. Our results show that the denoised single-repetition data are equivalent to ∼2.5-repetition averaged data in terms of image similarity quantified by MAD, PSNR and SSIM (Fig. 1), while being equivalent to 1.55 to 2.24-repetition averaged data in terms of cortical surface positioning and thickness estimation accuracy (Fig. 11). The discrepancy between the efficiency factors calculated using image and surface metrics suggests that the commonly used image similarity metrics alone cannot faithfully reflect the cortical surface reconstruction quality and the objective functions used during the denoising process, e.g., the mean squared error, can be further optimized for cortical surface reconstruction applications.

In addition to improving cortical surface reconstruction at sub-millimeter resolution, denoising methods can be also useful for improving cortical surface reconstruction at standard 1-mm isotropic resolution. The ∼5 minutes scan time of a standard 1-mm isotropic resolution acquisition (assuming a parallel imaging acceleration factor of two) is still considered long for elderly subjects, children and some patient populations prone to motion and/or discomfort. In order to save scan time, T_1_-weighted images at resolutions lower than 1-mm isotropic (e.g., 3 to 5-mm axial slices in a 2-dimensional acquisition, ∼1.5 mm resolution along the phase-encoding direction in a 3-dimensional acquisition) are also routinely acquired in clinical practice and large-scale neuroimaging studies such as the Parkinson Progression Marker Initiative^120^, which preclude accurate cortical surface reconstruction and quantitative analysis of cortical morphology at 1-mm isotropic resolution. In these applications, higher parallel imaging factors (e.g., 3 or 4) can be adopted for further shortening the 1-mm isotropic acquisition duration, coupled with denoising methods for accurate cortical surface reconstruction from the noisier images. More advanced fast imaging technologies such as Wave-CAIPI^53^ can achieve even higher acceleration factors (e.g., 9) with minimal noise and artifact penalties and have been shown to reduce the scan time of a 1-mm isotropic acquisition to 1.15 minutes with estimated regional brain tissue volumes comparable to those from a standard 5.2 minutes scan^121^. Denoising methods are expected to further improve the image quality of highly accelerated 1-mm isotropic data, acquired even within a minute, for accurately delineating the cortical surface^119^.

The pros and cons of different denoising methods need to be weighed for different applications. The three denoising methods evaluated here, DnCNN, BM4D and AONLM, are three well-known classical denoising methods with widely accepted superior denoising performance and are often used as references against which newly developed denoising methods are compared. Our results show that CNN-based denoising method DnCNN generates denoised images and consequently reconstructed cortical surfaces that are more similar to the ground truth than BM4D and AONLM, which is achieved by effectively exploiting the data redundancy in local and non-local spatial locations as well as across numerous subjects. However, the data-driven nature of DnCNN requires additional training data (e.g., data from 4 subjects in our study) for optimizing parameters or fine-tuning the parameters of an existing DnCNN pre-trained on other datasets for improved generalization across hardware systems, pulse sequences and sequences parameters. It has been shown that results obtained by directly applying CNNs pre-trained on a different dataset are reasonably accurate but are slightly worse compared to those obtained from fine-tuned CNNs^47,122,123^. On the other hand, BM4D and AONLM only require a single input noisy image volume, which are more convenient in practice and especially useful when the training data for DnCNN are not available (e.g., denoising legacy data). In terms of computational efficiency, DnCNN, once trained, can be executed in milliseconds or seconds, much faster than BM4D and AONLM with run times that last anywhere from minutes to an hour depending on the computing platform. The extremely fast DnCNN could facilitate review of denoised images in near real-time on the host computer of the MRI console.

The specific DnCNN network we chose for our study is a pioneering, classic CNN for noise removal, which is characterized by its simplest (i.e., cascaded convolutional filters and nonlinear activation functions) yet very deep (20 layers) network topology. The “plain” network architecture achieves superior performance not only in noise removal^77^, but also vision tasks such as object recognition^124^ and super-resolution^109,125^. DnCNN has been demonstrated to outperform the state-of-the-art BM3D denoising for 2-dimension natural images, which is reproduced on empirical 3-dimenstion MRI data in our study in terms of both image quality and cortical surface positioning. The versatile DnCNN can be also used to perform simultaneous denoising and super-resolution (referred to as VDSR in the context of super-resolution^109^). This versatility is beneficial for obtaining high-quality images at a true, high sub-millimeter resolution using only product sequences and reconstruction provided by vendors, which can only encode a whole-brain volume at a high resolution nominally because of the use of partial Fourier imaging and the increased T_1_ blurring during the inversion recovery. Simultaneous denoising and super-resolution is also helpful for obtaining images at ultra-high resolution (e.g., 0.25-mm isotropic resolution^126^, which requires averaging 8 repetitions from 10 hour scan on a 7-Tesla scanner) by making a balance between extremely challenging individual denoising or super-resolution and avoiding the use of prospective motion correction to acquire input noisy images at native ultra-high resolution for denoising^127–131^. Image super-resolution has also been shown to be an alternative way to obtain high-quality sub-millimeter resolution images for improving cortical surface reconstruction^47^.

## Conclusion

Our unique dataset and systematic characterization of different denoising methods support the use of denoising methods for improving cortical surface reconstruction at sub-millimeter resolution from single-repetition anatomical MR images. The denoised images from three classical denoising methods, namely DnCNN, BM4D and AONLM, are shown to be similar to 6-repetition averaged images, with high image similarity quantified by MAD, PSNR and SSIM that are equivalent to those of T_1_-weighted images obtained by averaging ∼2.5 repetitions of the data. The discrepancies in the gray–white surface placement, gray–CSF surface placement and cortical thickness estimation are lower than 165 μm, equivalent to that of T_1_-weighted images obtained by averaging ∼1.55 to 2.24 repetitions of the data and on the same order as the test-retest precision of FreeSurfer. The scan-rescan precision of the cortical surface positioning and thickness estimation from the denoised single-repetition data is lower than 170 μm. The proposed approach would facilitate the adoption of next-generation cortical surface reconstruction methods in a wider range of applications and neuroimaging studies that require accurate models of cortical anatomy.

## Acknowledgments

Thanks to Mr. Ned Ohringer for assistance in volunteer recruitment and data collection. We gratefully acknowledge the use of the Siemens “Works-In-Progress” package WIP900, which provided the inner-loop accelerated MPRAGE pulse sequence and image reconstruction. This work was supported by the National Institutes of Health (grant numbers P41-EB015896, P41-EB030006, R01-EB023281, K23-NS096056, R01-MH111419, R01-MH124004, U01-EB026996, S10-RR023401, S10-RR019307 and S10-RR023043), a Massachusetts General Hospital Claflin Distinguished Scholar Award, and the Athinoula A. Martinos Center for Biomedical Imaging. B.B. has provided consulting services to Subtle Medical.

## Data and code availability

The source code of the DnCNN network implemented using Keras API and the pre-trained models using the MGH 0.6-mm isotropic data will be available at https://github.com/qiyuantian/DnSurfer.

## Supplementary Information

**Supplementary Figure 1.**
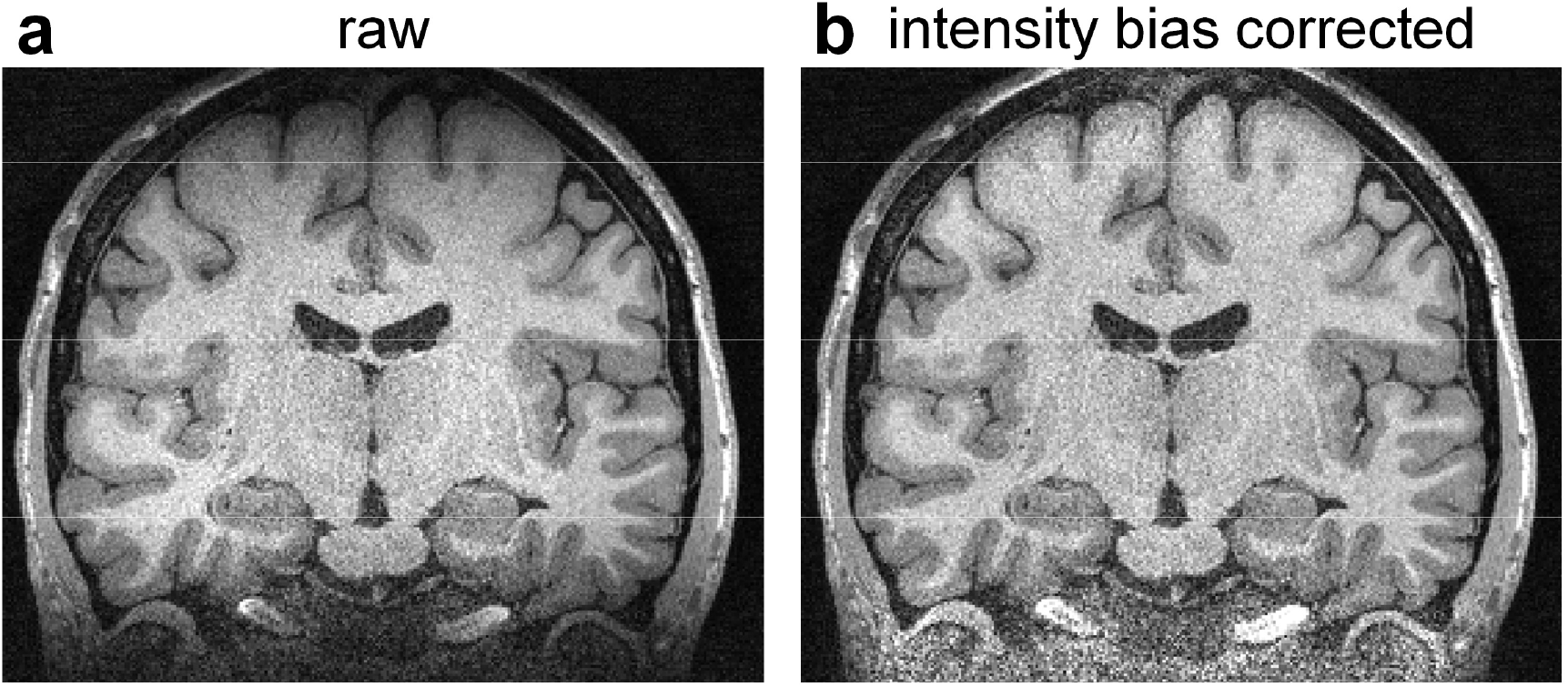
Intensity bias. A representative coronal image slice from the raw data acquired using a slab-selective oblique-axial acquisition (a) and the same image slice corrected for the spatially varying intensity bias induced by the B1 field inhomogeneity (b).

**Supplementary Figure 2.**
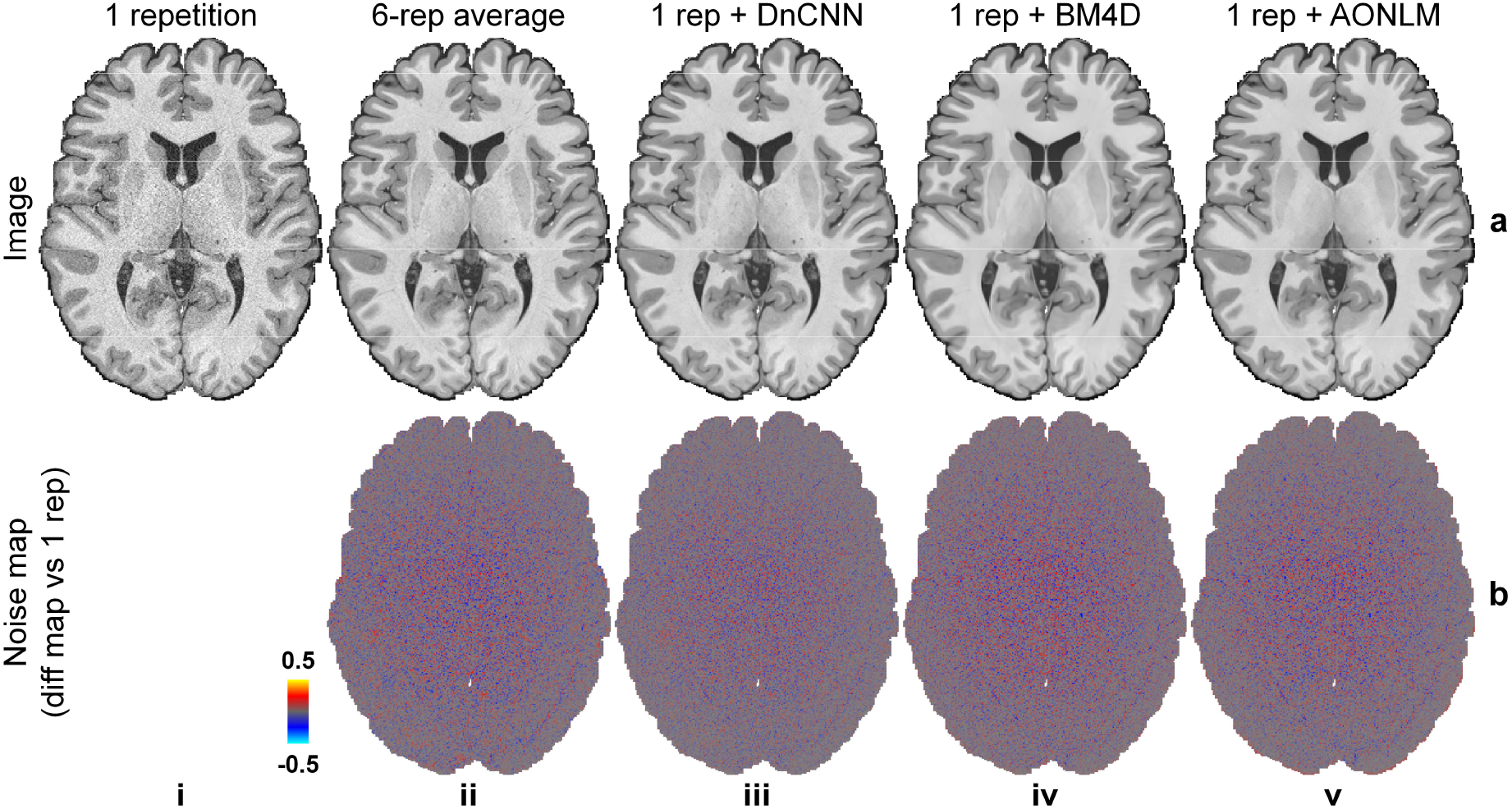
Noise estimation. A representative axial image slice from the single-repetition data (a, i), the 6-repetition averaged data (a, ii), and the single-repetition data denoised by DnCNN (a, iii), BM4D (a, iv) and AONLM (a, v), and their residuals (i.e., estimated noise) compared to the image from the single-repetition data (rows a, i).

**Supplementary Figure 3.**
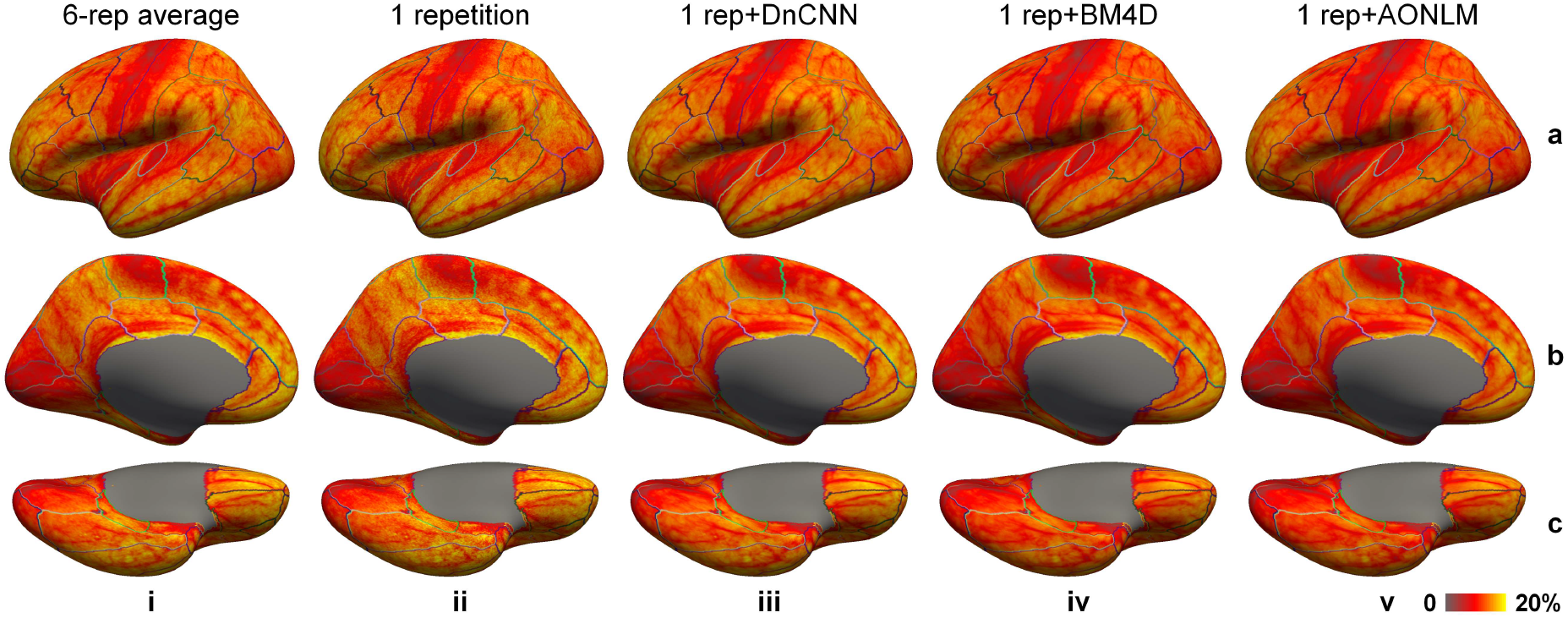
Averaged gray–white contrast. Left-hemisphere vertex-wise contrast between the gray matter and white matter image intensity (expressed as [white − gray]/[white + gray]⋅100%) from the 6-repetition averaged data (column i), single-repetition data (column ii), single-repetition data denoised by DnCNN (column iii), BM4D (column iv) and AONLM (column v) averaged across 30 image volumes from the 5 evaluation subjects, displayed on inflated surface representations. Different cortical regions from the FreeSurfer cortical parcellation (i.e., *aparc.annot*) are depicted as colored outlines.

**Supplementary Figure 4.**
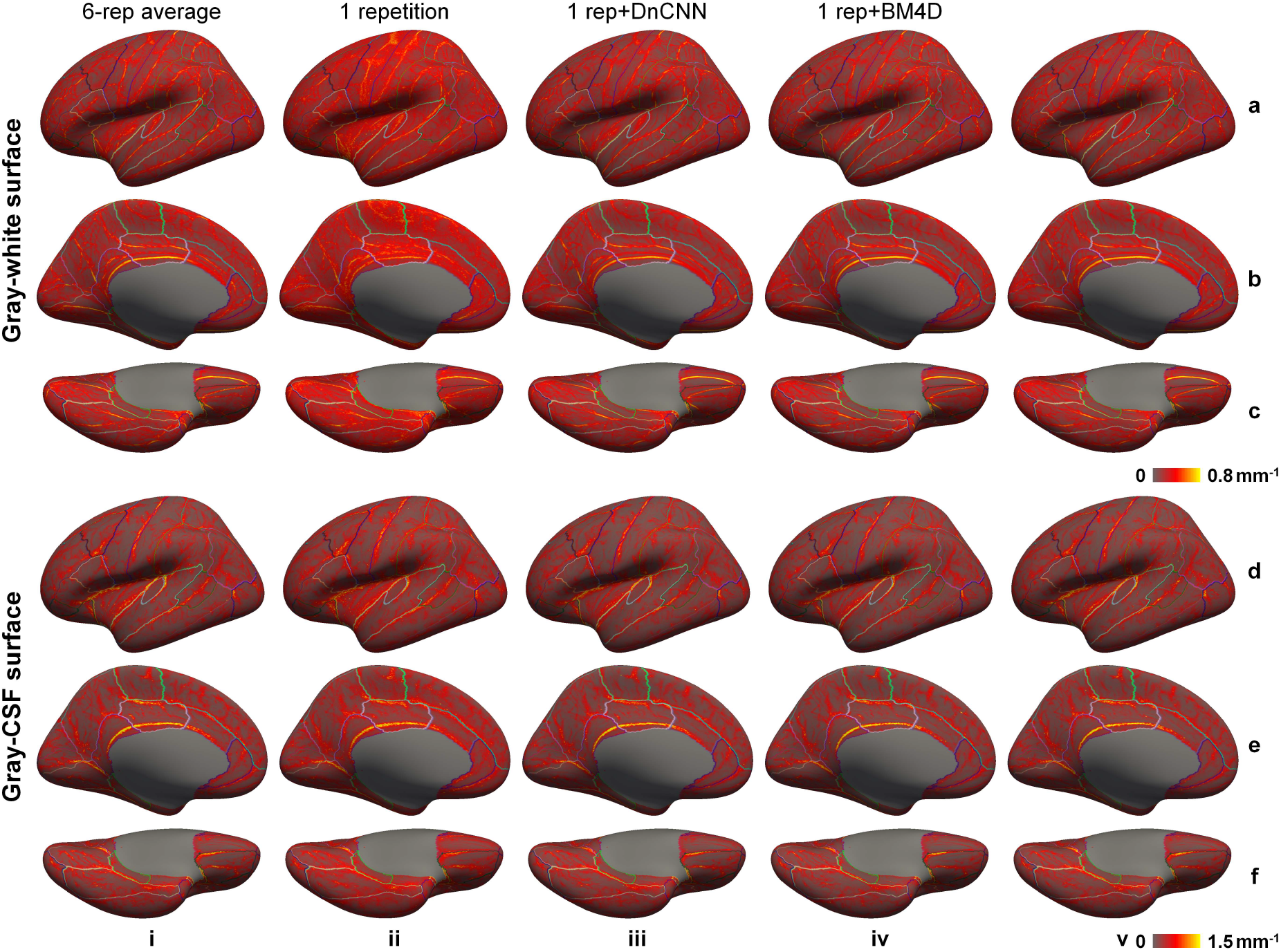
Averaged cortical surface smoothness. Left-hemisphere vertex-wise mean curvature of the reconstructed gray–white surfaces (a–c) and gray–CSF surfaces (d–f) from the 6-repetition averaged data (column i), single-repetition data (column ii), single-repetition data denoised by DnCNN (column iii), BM4D (column iv) and AONLM (column v) averaged across 30 image volumes from the 5 evaluation subjects, displayed on inflated surface representations. Different cortical regions from the FreeSurfer cortical parcellation (i.e., *aparc.annot*) are depicted as colored outlines.

**Supplementary Table 1.** Gray–white surface displacement. The group mean and standard deviation of the mean absolute displacement of the gray–white surface estimated from the images from the single-repetition data, 2- to 5-repetition averaged data and the single-repetition data denoised by DnCNN, BM4D and AONLM compared to the images from the 6-repetition averaged data across 30 image volumes from the 5 evaluation subjects, calculated for 34 cortical parcels (left and right hemispheres combined) from the Desikan-Killiany Atlas provided by FreeSurfer and for the whole brain.

| Gray-white surface displacement (mm) compared to ground truth from 6-repetition averaged data |  |  |  |  |  |  |  |  |  |
| --- | --- | --- | --- | --- | --- | --- | --- | --- | --- |
| # | Cortices | 1 repetition | 2 repetitions | 3 repetitions | 4 repetitions | 5 repetitions | 1 rep + DnCNN | 1 rep + BM4D | 1 rep + AONLM |
| 1 | Superior frontal | 0.2724±0.0536 | 0.1692±0.03157 | 0.129±0.01805 | 0.1082±0.01406 | 0.09206±0.01484 | 0.1485±0.02164 | 0.168±0.02684 | 0.1669±0.02786 |
| 2 | Rostral middle frontal | 0.1977±0.07068 | 0.1252±0.03976 | 0.09743±0.02443 | 0.0831±0.02133 | 0.07218±0.02121 | 0.1172±0.02972 | 0.1311±0.03322 | 0.1271±0.03226 |
| 3 | Caudal middle frontal | 0.2647±0.06135 | 0.1636±0.03242 | 0.1211±0.01899 | 0.09885±0.01269 | 0.08617±0.01592 | 0.1397±0.01954 | 0.1533±0.02307 | 0.1531±0.02558 |
| 4 | Pars opercularis | 0.2544±0.07431 | 0.1517±0.03902 | 0.11±0.02165 | 0.09073±0.01774 | 0.077±0.01745 | 0.1235±0.02539 | 0.1353±0.02765 | 0.1352±0.02928 |
| 5 | Pars triangularis | 0.2372±0.1074 | 0.1502±0.06582 | 0.1069±0.03371 | 0.08975±0.02797 | 0.07695±0.02918 | 0.1206±0.03325 | 0.133±0.03641 | 0.1302±0.03755 |
| 6 | Pars orbitalis | 0.2248±0.08882 | 0.1518±0.05481 | 0.1165±0.0354 | 0.09857±0.03102 | 0.08235±0.02642 | 0.1425±0.04126 | 0.1535±0.04307 | 0.1489±0.04144 |
| 7 | Lateral orbitofrontal | 0.2732±0.06782 | 0.1841±0.04273 | 0.1381±0.02464 | 0.1175±0.02447 | 0.09621±0.01819 | 0.1544±0.03255 | 0.1691±0.03208 | 0.1717±0.03445 |
| 8 | Medial orbitofrontal | 0.2302±0.0599 | 0.1605±0.03942 | 0.13±0.0301 | 0.1101±0.02623 | 0.09281±0.02129 | 0.1492±0.03357 | 0.157±0.03212 | 0.1587±0.03261 |
| 9 | Precentral | 0.33±0.05801 | 0.2106±0.0352 | 0.1655±0.02341 | 0.1374±0.01881 | 0.1204±0.0205 | 0.1835±0.03179 | 0.2068±0.04857 | 0.2135±0.0479 |
| 10 | Paracentral | 0.3474±0.05652 | 0.2146±0.03067 | 0.1687±0.02053 | 0.1419±0.01826 | 0.1231±0.02151 | 0.1905±0.02711 | 0.2162±0.05089 | 0.2287±0.05571 |
| 11 | Frontal pole | 0.2304±0.0875 | 0.1612±0.05737 | 0.1396±0.04845 | 0.1154±0.03685 | 0.107±0.04082 | 0.1659±0.06649 | 0.194±0.07748 | 0.1794±0.07227 |
| 12 | Superior parietal | 0.2165±0.04414 | 0.1303±0.01554 | 0.1018±0.009543 | 0.08448±0.006521 | 0.07332±0.009185 | 0.1235±0.01562 | 0.1387±0.02022 | 0.1396±0.01839 |
| 13 | Inferior parietal | 0.183±0.03884 | 0.1148±0.01074 | 0.09062±0.006357 | 0.07635±0.005439 | 0.06773±0.008305 | 0.1125±0.01452 | 0.1255±0.0205 | 0.1232±0.0188 |
| 14 | Supramarginal | 0.1982±0.0372 | 0.1247±0.02024 | 0.09831±0.01181 | 0.08271±0.01015 | 0.07243±0.01124 | 0.1194±0.01627 | 0.1315±0.02106 | 0.1294±0.02088 |
| 15 | Postcentral | 0.2642±0.04854 | 0.1696±0.03413 | 0.1317±0.0239 | 0.1108±0.02018 | 0.09446±0.01912 | 0.1592±0.02723 | 0.1811±0.03708 | 0.1821±0.03282 |
| 16 | Precuneus | 0.2193±0.04294 | 0.1307±0.01409 | 0.1022±0.008607 | 0.0828±0.005642 | 0.07241±0.008587 | 0.1215±0.01296 | 0.1371±0.01762 | 0.1411±0.01781 |
| 17 | Superior temporal | 0.2593±0.03898 | 0.1726±0.02397 | 0.1315±0.01443 | 0.1144±0.01525 | 0.09869±0.01894 | 0.1555±0.01925 | 0.1662±0.0186 | 0.1665±0.01921 |
| 18 | Middle temporal | 0.2215±0.03958 | 0.144±0.02022 | 0.111±0.01282 | 0.09557±0.01214 | 0.08252±0.0128 | 0.1375±0.0191 | 0.1496±0.02422 | 0.1473±0.02215 |
| 19 | Inferior temporal | 0.2412±0.04746 | 0.1554±0.02683 | 0.122±0.01598 | 0.1045±0.01665 | 0.08684±0.01378 | 0.1526±0.02744 | 0.1618±0.02542 | 0.1643±0.02768 |
| 20 | Banks of sup temp sulcus | 0.1894±0.03193 | 0.1153±0.0147 | 0.08612±0.005842 | 0.07122±0.005444 | 0.05917±0.006335 | 0.09891±0.01159 | 0.1115±0.01577 | 0.1099±0.01658 |
| 21 | Fusiform | 0.2463±0.05098 | 0.1546±0.03042 | 0.1196±0.01901 | 0.09856±0.014 | 0.08318±0.01684 | 0.1451±0.02223 | 0.1566±0.0217 | 0.1619±0.02429 |
| 22 | Transverse temporal | 0.3646±0.07071 | 0.2309±0.05088 | 0.1799±0.03548 | 0.1396±0.03033 | 0.1224±0.02988 | 0.204±0.0425 | 0.2192±0.05211 | 0.2423±0.04297 |
| 23 | Entorhinal | 0.3434±0.06467 | 0.2577±0.08661 | 0.2046±0.04682 | 0.1725±0.03008 | 0.1477±0.02886 | 0.2442±0.04564 | 0.2839±0.05642 | 0.2853±0.06142 |
| 24 | Temporal pole | 0.3374±0.06953 | 0.252±0.06244 | 0.2046±0.0486 | 0.1732±0.03477 | 0.143±0.02987 | 0.2444±0.05361 | 0.3052±0.07984 | 0.3003±0.07335 |
| 25 | Parahippocampal | 0.2913±0.05749 | 0.192±0.03799 | 0.1455±0.01961 | 0.1266±0.01905 | 0.1086±0.01845 | 0.167±0.02443 | 0.1936±0.02066 | 0.1936±0.0291 |
| 26 | Insula | 0.3808±0.05685 | 0.2648±0.03606 | 0.2082±0.02442 | 0.1774±0.0183 | 0.1555±0.02039 | 0.2568±0.02525 | 0.3028±0.03278 | 0.3158±0.03693 |
| 27 | Lateral occipital | 0.1969±0.03237 | 0.1251±0.01119 | 0.1016±0.008153 | 0.08696±0.008086 | 0.07763±0.007071 | 0.1489±0.0266 | 0.1594±0.02523 | 0.1505±0.02395 |
| 28 | Lingual | 0.2352±0.03415 | 0.1624±0.01964 | 0.1348±0.01657 | 0.1168±0.01676 | 0.103±0.01289 | 0.2157±0.03783 | 0.2265±0.04672 | 0.2282±0.0414 |
| 29 | Cuneus | 0.2169±0.04204 | 0.1433±0.01586 | 0.122±0.01576 | 0.1037±0.0135 | 0.09455±0.01412 | 0.1791±0.03972 | 0.189±0.03014 | 0.1797±0.02911 |
| 30 | Pericalcarine | 0.2432±0.03715 | 0.1708±0.02007 | 0.1444±0.02321 | 0.1285±0.02209 | 0.1181±0.02442 | 0.2444±0.05763 | 0.2452±0.0581 | 0.2351±0.04442 |
| 31 | Rostral anterior cingulate | 0.3113±0.07176 | 0.1987±0.04196 | 0.1517±0.03522 | 0.1219±0.02441 | 0.1028±0.01943 | 0.1596±0.03249 | 0.1675±0.03063 | 0.1692±0.02938 |
| 32 | Caudal anterior cingulate | 0.2847±0.05613 | 0.1869±0.0353 | 0.1376±0.01828 | 0.1121±0.01195 | 0.09756±0.01809 | 0.1487±0.02033 | 0.1574±0.02267 | 0.1572±0.02259 |
| 33 | Posterior cingulate | 0.3±0.04635 | 0.1798±0.03109 | 0.1306±0.01324 | 0.1064±0.01108 | 0.09014±0.01407 | 0.1452±0.02111 | 0.1552±0.0195 | 0.1604±0.01883 |
| 34 | Isthmus cingulate | 0.3189±0.07067 | 0.2002±0.03621 | 0.1539±0.02359 | 0.127±0.01811 | 0.1099±0.02011 | 0.1734±0.02555 | 0.1886±0.0248 | 0.1925±0.02475 |
| 35 | Whole brain | 0.2471±0.04139 | 0.1578±0.02556 | 0.1229±0.01523 | 0.1033±0.01237 | 0.08925±0.01333 | 0.1493±0.02013 | 0.1647±0.02477 | 0.165±0.02452 |

**Supplementary Table 2.**
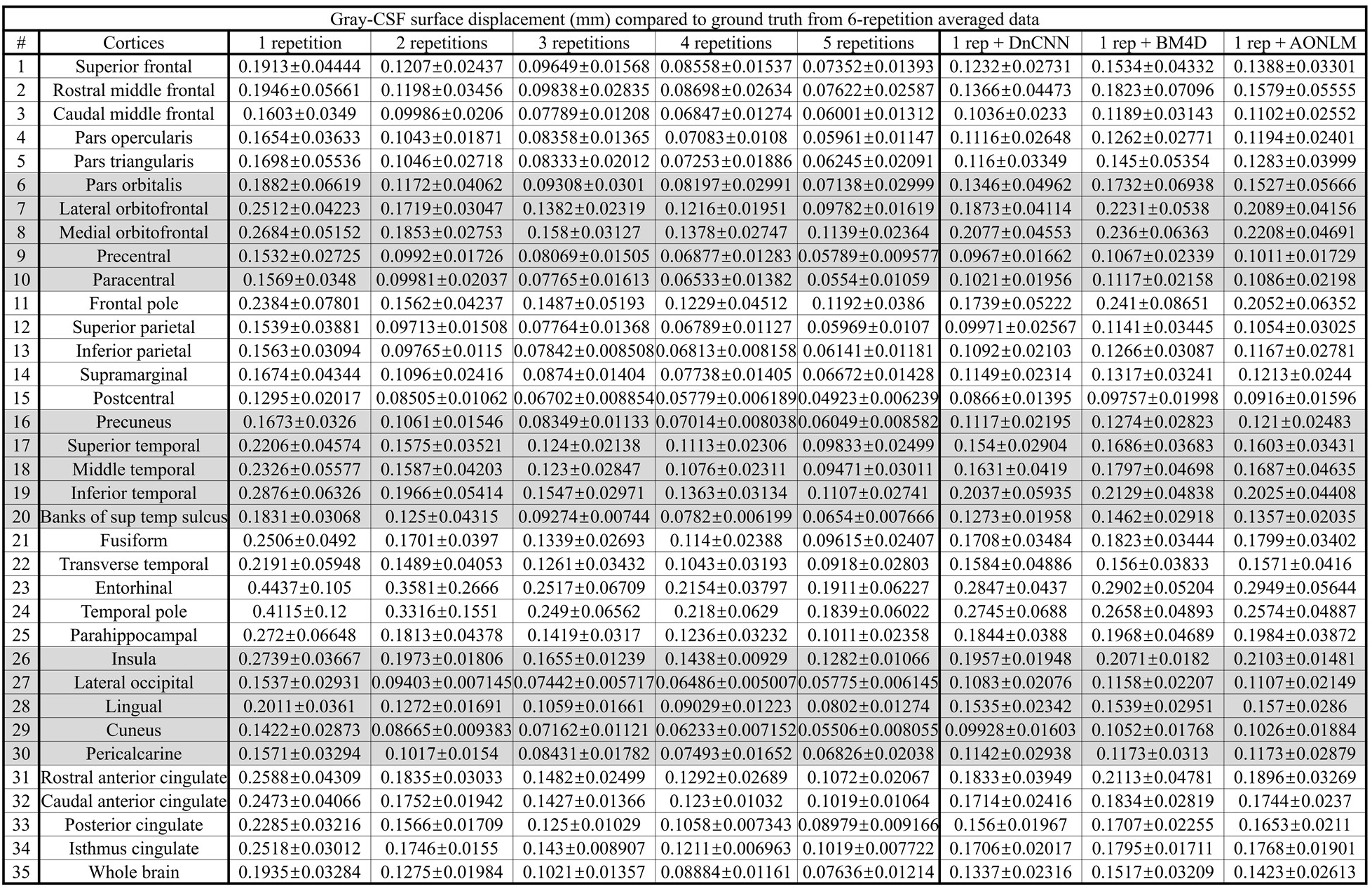
Gray–CSF surface displacement. The group mean and standard deviation of the mean absolute displacement of the gray–CSF surface estimated from the images from the single-repetition data, 2- to 5-repetition averaged data and the single-repetition data denoised by DnCNN, BM4D and AONLM compared to the images from the 6-repetition averaged data across 30 image volumes from the 5 evaluation subjects, calculated for 34 cortical parcels (left and right hemispheres combined) from the Desikan-Killiany Atlas provided by FreeSurfer and for the whole brain.

**Supplementary Table 3.** Cortical thickness difference. The group mean and standard deviation of the mean absolute difference of the cortical thickness estimated from the images from the single-repetition data, 2- to 5-repetition averaged data and the single-repetition data denoised by DnCNN, BM4D and AONLM compared to the images from the 6-repetition averaged data across 30 image volumes from the 5 evaluation subjects, calculated for 34 cortical parcels (left and right hemispheres combined) from the Desikan-Killiany Atlas provided by FreeSurfer and for the whole brain. displacement/difference of the gray–white surface, gray–CSF surface and cortical thickness estimated from two consecutively acquired 1-repetition data denoised by DnCNN, BM4D and AONLM across 25 image volumes from the 5 evaluation subjects, calculated for 34 cortical parcels (left and right hemispheres combined) from the Desikan-Killiany Atlas provided by FreeSurfer and for the whole brain.

| Cortical thickness difference (mm) compared to ground truth from 6-repetition averaged data |  |  |  |  |  |  |  |  |  |
| --- | --- | --- | --- | --- | --- | --- | --- | --- | --- |
| # | Cortices | 1 repetition | 2 repetitions | 3 repetitions | 4 repetitions | 5 repetitions | 1 rep + DnCNN | 1 rep + BM4D | 1 rep + AONLM |
| 1 | Superior frontal | 0.1853±0.03055 | 0.1186±0.02105 | 0.09382±0.01344 | 0.08042±0.01117 | 0.06817±0.01079 | 0.1214±0.02253 | 0.1499±0.03158 | 0.1373±0.02657 |
| 2 | Rostral middle frontal | 0.1631±0.04248 | 0.1009±0.02796 | 0.08117±0.02128 | 0.06981±0.01771 | 0.05943±0.01533 | 0.1121±0.03357 | 0.1384±0.04659 | 0.1253±0.03917 |
| 3 | Caudal middle frontal | 0.1748±0.03459 | 0.1116±0.02328 | 0.08574±0.01335 | 0.07274±0.01203 | 0.06092±0.01087 | 0.1112±0.02092 | 0.1303±0.02641 | 0.1239±0.02395 |
| 4 | Pars opercularis | 0.1714±0.0379 | 0.1063±0.02142 | 0.08087±0.01459 | 0.06865±0.0119 | 0.05729±0.009922 | 0.107±0.02624 | 0.1203±0.02687 | 0.116±0.02429 |
| 5 | Pars triangularis | 0.172±0.06022 | 0.1079±0.0393 | 0.08104±0.02311 | 0.06843±0.01959 | 0.05813±0.01807 | 0.1077±0.03285 | 0.1289±0.04786 | 0.1196±0.03938 |
| 6 | Pars orbitalis | 0.1683±0.04852 | 0.1111±0.03545 | 0.0875±0.0284 | 0.07326±0.02176 | 0.06187±0.01856 | 0.1207±0.03792 | 0.1425±0.05234 | 0.1306±0.0429 |
| 7 | Lateral orbitofrontal | 0.203±0.03651 | 0.141±0.02708 | 0.112±0.01909 | 0.09671±0.01592 | 0.0787±0.01252 | 0.1514±0.03095 | 0.1775±0.04205 | 0.1705±0.03593 |
| 8 | Medial orbitofrontal | 0.2003±0.03826 | 0.1416±0.02494 | 0.121±0.02194 | 0.1031±0.0182 | 0.08639±0.01666 | 0.1628±0.03482 | 0.18±0.04679 | 0.1752±0.03646 |
| 9 | Precentral | 0.1922±0.03063 | 0.1264±0.01884 | 0.1014±0.01442 | 0.08396±0.011 | 0.07186±0.01153 | 0.1325±0.02406 | 0.1493±0.03568 | 0.1528±0.03431 |
| 10 | Paracentral | 0.1986±0.0269 | 0.1295±0.01957 | 0.1025±0.01324 | 0.08482±0.01074 | 0.07088±0.01206 | 0.1395±0.02098 | 0.1539±0.03789 | 0.1625±0.03939 |
| 11 | Frontal pole | 0.1861±0.05158 | 0.1253±0.03208 | 0.1114±0.03126 | 0.09341±0.02309 | 0.08576±0.01941 | 0.1496±0.04203 | 0.1839±0.06527 | 0.1615±0.04754 |
| 12 | Superior parietal | 0.1491±0.02225 | 0.09529±0.01024 | 0.07489±0.006546 | 0.06361±0.004999 | 0.05417±0.005375 | 0.1041±0.02195 | 0.1183±0.02581 | 0.1142±0.02263 |
| 13 | Inferior parietal | 0.1397±0.02422 | 0.08831±0.009968 | 0.07041±0.005213 | 0.06014±0.005024 | 0.05274±0.005975 | 0.0965±0.01778 | 0.1119±0.02005 | 0.1066±0.02054 |
| 14 | Supramarginal | 0.1485±0.03003 | 0.09654±0.01706 | 0.07699±0.01058 | 0.0663±0.008693 | 0.05666±0.008479 | 0.1019±0.01817 | 0.1159±0.0227 | 0.1096±0.01876 |
| 15 | Postcentral | 0.1665±0.02588 | 0.1092±0.01778 | 0.08545±0.01273 | 0.07156±0.01021 | 0.05974±0.009252 | 0.1175±0.02093 | 0.1335±0.02838 | 0.1337±0.02467 |
| 16 | Precuneus | 0.1488±0.02005 | 0.09779±0.01061 | 0.07796±0.007576 | 0.06417±0.005039 | 0.05462±0.005488 | 0.1044±0.01512 | 0.1199±0.01846 | 0.119±0.01758 |
| 17 | Superior temporal | 0.1912±0.02614 | 0.133±0.02027 | 0.1047±0.01179 | 0.09284±0.01161 | 0.07931±0.01318 | 0.1332±0.01916 | 0.1487±0.02502 | 0.1425±0.02208 |
| 18 | Middle temporal | 0.1845±0.03134 | 0.1273±0.02497 | 0.1014±0.01732 | 0.08789±0.01388 | 0.07544±0.01631 | 0.1346±0.02815 | 0.1538±0.03099 | 0.1462±0.02955 |
| 19 | Inferior temporal | 0.2074±0.04201 | 0.1461±0.03331 | 0.1166±0.01991 | 0.1016±0.02024 | 0.08295±0.01691 | 0.1599±0.04164 | 0.1738±0.03635 | 0.1646±0.03358 |
| 20 | Banks of sup temp sulcus | 0.155±0.02527 | 0.1023±0.0263 | 0.0763±0.00614 | 0.06423±0.004637 | 0.0536±0.005343 | 0.1055±0.01555 | 0.1218±0.02029 | 0.1139±0.01642 |
| 21 | Fusiform | 0.1926±0.03602 | 0.136±0.02807 | 0.1069±0.01987 | 0.09074±0.01791 | 0.07546±0.01684 | 0.1387±0.02418 | 0.1572±0.0279 | 0.155±0.02842 |
| 22 | Transverse temporal | 0.2331±0.0421 | 0.1492±0.03417 | 0.1154±0.02583 | 0.0932±0.02032 | 0.08142±0.01675 | 0.1483±0.02395 | 0.154±0.02554 | 0.1657±0.02443 |
| 23 | Entorhinal | 0.3209±0.07421 | 0.2632±0.1661 | 0.1881±0.05271 | 0.1585±0.02637 | 0.1416±0.03688 | 0.2265±0.04005 | 0.2367±0.05287 | 0.2367±0.05558 |
| 24 | Temporal pole | 0.2957±0.08222 | 0.2418±0.1101 | 0.1869±0.04348 | 0.1599±0.03715 | 0.1379±0.03905 | 0.2201±0.05038 | 0.2372±0.05409 | 0.2288±0.05605 |
| 25 | Parahippocampal | 0.2255±0.03986 | 0.1611±0.03048 | 0.1259±0.02137 | 0.1108±0.02247 | 0.09077±0.0186 | 0.1687±0.03112 | 0.1859±0.03972 | 0.1829±0.03407 |
| 26 | Insula | 0.242±0.0318 | 0.1763±0.01763 | 0.1456±0.01314 | 0.126±0.01032 | 0.1136±0.01147 | 0.182±0.01983 | 0.2041±0.02418 | 0.2057±0.02263 |
| 27 | Lateral occipital | 0.1447±0.02396 | 0.09005±0.006984 | 0.07283±0.003932 | 0.06242±0.003198 | 0.05505±0.00378 | 0.1089±0.01737 | 0.1212±0.01866 | 0.1153±0.01893 |
| 28 | Lingual | 0.1703±0.02699 | 0.1165±0.01274 | 0.09657±0.01102 | 0.08212±0.008749 | 0.07114±0.007885 | 0.1557±0.02316 | 0.1648±0.03295 | 0.166±0.0262 |
| 29 | Cuneus | 0.1442±0.02413 | 0.09269±0.00823 | 0.07796±0.00646 | 0.06602±0.005539 | 0.05728±0.005062 | 0.118±0.02291 | 0.1261±0.01881 | 0.1222±0.02001 |
| 30 | Pericalcarine | 0.1617±0.02855 | 0.1107±0.01377 | 0.09247±0.01198 | 0.08154±0.01136 | 0.07259±0.01324 | 0.1669±0.04166 | 0.1619±0.04207 | 0.1551±0.03643 |
| 31 | Rostral anterior cingulate | 0.214±0.03345 | 0.1449±0.02209 | 0.1152±0.01989 | 0.09787±0.01866 | 0.08315±0.01403 | 0.146±0.03294 | 0.1647±0.03866 | 0.1539±0.03146 |
| 32 | Caudal anterior cingulate | 0.1982±0.02901 | 0.1408±0.01927 | 0.1143±0.01261 | 0.097±0.009672 | 0.08374±0.01348 | 0.1422±0.01745 | 0.1507±0.02224 | 0.1455±0.02152 |
| 33 | Posterior cingulate | 0.1947±0.02562 | 0.1313±0.01813 | 0.1034±0.01065 | 0.08557±0.007924 | 0.0729±0.008893 | 0.1345±0.0187 | 0.1433±0.02052 | 0.1449±0.02004 |
| 34 | Isthmus cingulate | 0.2176±0.03474 | 0.1465±0.01827 | 0.1189±0.01218 | 0.1007±0.009819 | 0.08355±0.01006 | 0.1493±0.01937 | 0.1559±0.01735 | 0.1571±0.02014 |
| 35 | Whole brain | 0.1761±0.02746 | 0.1171±0.01801 | 0.09334±0.01196 | 0.07959±0.009615 | 0.06773±0.00934 | 0.1261±0.02026 | 0.1434±0.02681 | 0.1383±0.02349 |

**Supplementary Table 4.** Scan-rescan precision. The group mean and standard deviation of the mean absolute displacement/difference of the gray–white surface, gray–CSF surface and cortical thickness estimated from two consecutively acquired 1-repetition data denoised by DnCNN, BM4D and AONLM across 25 image volumes from the 5 evaluation subjects, calculated for 34 cortical parcels (left and right hemispheres combined) from the Desikan-Killiany Atlas provided by FreeSurfer and for the whole brain.

| Scan-rescan precision |  |  |  |  |  |  |  |  |  |  |
| --- | --- | --- | --- | --- | --- | --- | --- | --- | --- | --- |
| # | Cortices | Gray-white surface displacement (mm) |  |  | Gray-CSF surface displacement (mm) |  |  | Cortical thickness difference (mm) |  |  |
|  |  | 1 rep + DnCNN | 1 rep + BM4D | 1 rep + AONLM | 1 rep + DnCNN | 1 rep + BM4D | 1 rep + AONLM | 1 rep + DnCNN | 1 rep + BM4D | 1 rep + AONLM |
| 1 | Superior frontal | 0.157±0.04002 | 0.159±0.04046 | 0.1659±0.04202 | 0.1518±0.04321 | 0.1703±0.04732 | 0.1665±0.0444 | 0.1452±0.03596 | 0.1608±0.03906 | 0.1595±0.03903 |
| 2 | Rostral middle frontal | 0.1425±0.05495 | 0.1592±0.05791 | 0.1621±0.05646 | 0.18±0.08167 | 0.2153±0.08891 | 0.2081±0.08159 | 0.1394±0.05284 | 0.1608±0.05442 | 0.1608±0.05269 |
| 3 | Caudal middle frontal | 0.147±0.03577 | 0.15±0.03873 | 0.1586±0.04249 | 0.1298±0.03596 | 0.138±0.04208 | 0.1403±0.03943 | 0.133±0.03158 | 0.1426±0.0358 | 0.1463±0.03562 |
| 4 | Pars opercularis | 0.1382±0.03913 | 0.1371±0.03691 | 0.1446±0.03969 | 0.1426±0.03819 | 0.143±0.03684 | 0.1445±0.03545 | 0.1339±0.03394 | 0.1376±0.03359 | 0.1405±0.03507 |
| 5 | Pars triangularis | 0.1403±0.05566 | 0.1504±0.06056 | 0.1524±0.05812 | 0.1505±0.05755 | 0.1657±0.06702 | 0.1635±0.06425 | 0.1323±0.0489 | 0.1451±0.05364 | 0.1473±0.05508 |
| 6 | Pars orbitalis | 0.1749±0.07372 | 0.1893±0.07797 | 0.1894±0.07512 | 0.188±0.09946 | 0.2115±0.09887 | 0.2058±0.099 | 0.1546±0.0669 | 0.1748±0.06852 | 0.175±0.0693 |
| 7 | Lateral orbitofrontal | 0.1937±0.0651 | 0.1964±0.0644 | 0.2015±0.06355 | 0.2553±0.09246 | 0.2681±0.09147 | 0.2669±0.08636 | 0.1949±0.05837 | 0.2052±0.05872 | 0.2077±0.05615 |
| 8 | Medial orbitofrontal | 0.1877±0.05595 | 0.1887±0.05701 | 0.1953±0.05423 | 0.2737±0.09121 | 0.2817±0.09183 | 0.2814±0.08638 | 0.206±0.05959 | 0.2134±0.06522 | 0.2158±0.05669 |
| 9 | Precentral | 0.2019±0.04555 | 0.1941±0.04627 | 0.2064±0.04806 | 0.119±0.02472 | 0.1164±0.0256 | 0.1203±0.02376 | 0.1599±0.03272 | 0.1592±0.03615 | 0.1689±0.03603 |
| 10 | Paracentral | 0.2125±0.03338 | 0.1972±0.03643 | 0.2208±0.04084 | 0.1288±0.02507 | 0.1211±0.02566 | 0.1269±0.02685 | 0.1713±0.02595 | 0.1665±0.03158 | 0.1825±0.03496 |
| 11 | Frontal pole | 0.2133±0.1181 | 0.2412±0.1165 | 0.2415±0.1251 | 0.2369±0.1384 | 0.2956±0.117 | 0.2827±0.1149 | 0.1784±0.07987 | 0.214±0.08191 | 0.2148±0.07765 |
| 12 | Superior parietal | 0.138±0.01979 | 0.1364±0.02123 | 0.1449±0.02286 | 0.1192±0.03514 | 0.1255±0.03961 | 0.1258±0.03965 | 0.1238±0.02437 | 0.1296±0.02686 | 0.1337±0.02628 |
| 13 | Inferior parietal | 0.1191±0.01762 | 0.1263±0.02141 | 0.1311±0.02203 | 0.1277±0.0292 | 0.1397±0.03787 | 0.1389±0.0334 | 0.114±0.01778 | 0.1266±0.02381 | 0.1272±0.02193 |
| 14 | Supramarginal | 0.1285±0.02556 | 0.1316±0.02738 | 0.1374±0.02837 | 0.1387±0.0382 | 0.147±0.04231 | 0.1424±0.03633 | 0.1223±0.02435 | 0.1325±0.02711 | 0.131±0.02619 |
| 15 | Postcentral | 0.178±0.0376 | 0.1724±0.03859 | 0.1853±0.03925 | 0.1098±0.02188 | 0.1094±0.02426 | 0.1118±0.02179 | 0.1435±0.0263 | 0.1437±0.02952 | 0.1514±0.02882 |
| 16 | Precuneus | 0.1399±0.01962 | 0.1377±0.0201 | 0.1452±0.02017 | 0.1392±0.02875 | 0.137±0.03378 | 0.1379±0.03176 | 0.1321±0.01891 | 0.1345±0.02144 | 0.1376±0.02212 |
| 17 | Superior temporal | 0.1646±0.02638 | 0.1581±0.02687 | 0.1671±0.03005 | 0.1856±0.04356 | 0.183±0.04722 | 0.1851±0.04347 | 0.1588±0.02904 | 0.1662±0.03557 | 0.1673±0.03314 |
| 18 | Middle temporal | 0.148±0.02896 | 0.1503±0.0298 | 0.1564±0.03065 | 0.1902±0.0642 | 0.2059±0.07408 | 0.1996±0.072 | 0.1608±0.03892 | 0.1791±0.04963 | 0.1742±0.04722 |
| 19 | Inferior temporal | 0.1779±0.04484 | 0.1708±0.0389 | 0.1786±0.04239 | 0.2528±0.08932 | 0.2593±0.09676 | 0.253±0.0968 | 0.1951±0.05706 | 0.2092±0.0642 | 0.2027±0.06386 |
| 20 | Banks of sup temp sulcus | 0.1115±0.01632 | 0.1165±0.0205 | 0.1211±0.01895 | 0.1571±0.02971 | 0.1636±0.0388 | 0.1589±0.02817 | 0.1313±0.01807 | 0.1418±0.0248 | 0.1381±0.01861 |
| 21 | Fusiform | 0.1705±0.03561 | 0.1617±0.0289 | 0.1715±0.03161 | 0.2182±0.06523 | 0.207±0.06214 | 0.2095±0.06291 | 0.1772±0.04459 | 0.1801±0.04654 | 0.1799±0.04528 |
| 22 | Transverse temporal | 0.2239±0.04902 | 0.2251±0.06411 | 0.2557±0.0556 | 0.1767±0.04511 | 0.1559±0.04721 | 0.1594±0.03628 | 0.1743±0.02977 | 0.1789±0.03816 | 0.1966±0.03589 |
| 23 | Entorhinal | 0.3109±0.07561 | 0.284±0.05143 | 0.2988±0.05291 | 0.3575±0.09264 | 0.3187±0.1033 | 0.3445±0.1153 | 0.2805±0.07655 | 0.2742±0.07404 | 0.2781±0.08825 |
| 24 | Temporal pole | 0.3031±0.09927 | 0.2907±0.1182 | 0.3064±0.1129 | 0.3292±0.1547 | 0.3054±0.1297 | 0.3276±0.1617 | 0.2688±0.1027 | 0.2647±0.1076 | 0.2886±0.1264 |
| 25 | Parahippocampal | 0.1944±0.03907 | 0.1615±0.02378 | 0.1791±0.02739 | 0.2344±0.0733 | 0.2121±0.06963 | 0.2259±0.07579 | 0.2095±0.05711 | 0.1997±0.05964 | 0.2081±0.06089 |
| 26 | Insula | 0.2779±0.05112 | 0.2631±0.04618 | 0.261±0.04645 | 0.232±0.03522 | 0.2065±0.02563 | 0.2061±0.02456 | 0.2103±0.03394 | 0.2022±0.03098 | 0.1987±0.02966 |
| 27 | Lateral occipital | 0.1511±0.02218 | 0.1621±0.02349 | 0.1639±0.02562 | 0.1233±0.02559 | 0.1264±0.02715 | 0.1288±0.02704 | 0.1248±0.01818 | 0.1349±0.02073 | 0.136±0.02157 |
| 28 | Lingual | 0.2055±0.02287 | 0.2136±0.02852 | 0.2243±0.03003 | 0.1664±0.02971 | 0.1559±0.02905 | 0.1641±0.0291 | 0.1681±0.02101 | 0.1742±0.02509 | 0.1803±0.02616 |
| 29 | Cuneus | 0.1719±0.02902 | 0.1855±0.03175 | 0.1874±0.03012 | 0.1098±0.02337 | 0.1108±0.02402 | 0.1148±0.02367 | 0.1286±0.01702 | 0.1395±0.02138 | 0.1403±0.01902 |
| 30 | Pericalcarine | 0.211±0.03117 | 0.2216±0.03522 | 0.2279±0.03896 | 0.1248±0.03283 | 0.1261±0.02955 | 0.1327±0.03491 | 0.1608±0.02608 | 0.1642±0.03376 | 0.1671±0.03185 |
| 31 | Rostral anterior cingulate | 0.1819±0.05255 | 0.1788±0.04467 | 0.1798±0.04353 | 0.2347±0.08022 | 0.2372±0.06655 | 0.2279±0.0626 | 0.1797±0.05297 | 0.1857±0.04439 | 0.1798±0.0465 |
| 32 | Caudal anterior cingulate | 0.1698±0.03529 | 0.1666±0.03504 | 0.172±0.03783 | 0.2103±0.03663 | 0.1981±0.03693 | 0.2012±0.03323 | 0.1702±0.02681 | 0.1711±0.03071 | 0.171±0.0303 |
| 33 | Posterior cingulate | 0.172±0.02972 | 0.161±0.02872 | 0.1705±0.03047 | 0.194±0.03335 | 0.1801±0.03517 | 0.1854±0.03289 | 0.1661±0.0294 | 0.1651±0.03293 | 0.1655±0.03138 |
| 34 | Isthmus cingulate | 0.1944±0.03594 | 0.1684±0.02745 | 0.1797±0.02981 | 0.1979±0.02761 | 0.174±0.02661 | 0.1855±0.02794 | 0.173±0.02684 | 0.1544±0.02564 | 0.1615±0.02692 |
| 35 | Whole brain | 0.1649±0.03371 | 0.1654±0.0336 | 0.1726±0.0348 | 0.1648±0.04283 | 0.1701±0.0447 | 0.1702±0.04257 | 0.1519±0.03158 | 0.16±0.03421 | 0.1619±0.0338 |

